# Human Amniotic Epithelial Cells Promote Chx10^−^/Pax6^+^ Müller Glia Subpopulation Reprogramming into Photoreceptor-like Cells

**DOI:** 10.1101/2024.02.01.578388

**Authors:** Hui Gao, Zhiyuan Yin, Xiaona Huang, Yuxiao Zeng, Ting Zou, A Luodan, Zhe Cha, Xuan Cheng, Lingling Ge, Jiahui Kang, Xi Lin, Hong Gong, Jing Xie, Xiaotang Fan, Haiwei Xu

## Abstract

Reprogramming Müller glia to regenerate neurons is a promising strategy for treating retinal degeneration, but whether Müller glia contain subpopulations with different regenerative fates remains unclear. Here, using single-cell RNA-seq analysis and Müller glia lineage-tracing mice with retinal degeneration, we reveal that Müller glia were heterogeneous and identify a specific Müller glial subpopulation (Chx10^−^/Pax6^+^) in healthy retinas that is activated and migrate to the outer nuclear layer (ONL) during photoreceptor degeneration. Transplantation of human amniotic epithelial cells (hAECs) facilitates the activation and extensive migration of the Chx10^−^/Pax6^+^ Müller glial subpopulation to the ONL, where they are reprogrammed into photoreceptor-like cells. Mechanistically, hAECs degrade the inhibitory extracellular matrix through regulating matrix metalloproteinases, which probably induces remodeling of the microenvironment of Müller glia and contributes to cell reprogramming. Consequently, hAEC transplantation improves visual function in rd10 mice. Our findings uncover a distinctive Müller glial subpopulation with the potential for reprogramming into photoreceptors.

## Introduction

Retinal degeneration (RD) diseases such as age-related macular degeneration, glaucoma, and retinitis pigmentosa are characterized by the progressive loss of retinal neurons^1–3^. As the central nervous system of adult mammals lacks regenerative capacity, RD is devastating and eventually develops into complete blindness^2^. Stem cell-based cell therapy offers tremendous promise for treating RD characterized by neuronal loss^4^. Among these therapies, direct reprogramming/transdifferentiation *in vivo* to replenish lost neurons is gaining momentum^5^.

Determining appropriate cell type is critical for implementing reprogramming *in vivo*. Supporting cells in situ, such as astrocytes and NG2 cells (also known as oligodendrocyte progenitor cells), are reprogrammed into various neuronal cell types in the brain^5,6^. In the retina, Müller glia (MG) are ideal candidates for endogenous stem cells because of their abundance and cell fate plasticity^7,8^. MG spontaneously regenerate all retinal neurons following various injuries in the zebrafish retina^7^. In mammals, MG usually undergo gliosis rather than reprogramming in response to retinal injury^7^. Notably, some studies have reported that overexpression or knockdown of critical genes by adeno-associated virus breaks the intrinsic barriers and achieves direct/indirect conversion of MG into retinal neurons^9–12^. However, some findings are controversial for various reasons, particularly due to the leaky expression of viral vectors ^13,14^. Reprogramming efficiency and long-term effectiveness of viral vectors require further confirmation and improvement. The direct in vivo delivery of transcription factors using viral vectors usually causes unexpected off-target effects, immune responses, and genotoxicity^6,15^.

In addition to gene manipulation, transplanting extrinsic stem cells, such as hematopoietic and bone marrow stem cells, induces MG reprogramming through cell–cell fusion^16–18^. We have attempted to treat RD using stem cell transplantation; however, the cell source, ethical limitations, immune rejection, and safety issues remain unresolved^19–21^. Human amniotic epithelial cells (hAECs) derived from pluripotent epiblasts have multiple advantages, including pluripotent stemness similar to that of embryonic stem cells, sufficient quantity, extremely low immunogenicity, no tumorigenicity and their use is not subjected to ethical limitations^22^. Indeed, hAECs have been used in clinical trials and show immense potential for treating neurodegenerative diseases, including Alzheimer’s, and Parkinson’s diseases^23–25^. In addition, hAECs have the unique capacity to regulate the extracellular microenvironment, especially the extracellular matrix (ECM), when compared with other stem cells. hAECs improve tissue repair and regeneration by regulating ECM remodeling during lung fibrosis and premature ovarian failure^26,27^. Previously, photoreceptors differentiated from hAECs were transplanted into the retina to improve visual function in RD rats^28^. Whether direct hAECs transplantation can protect retinal function and promote MG reprogramming has not been elucidated.

Previous studies have shown that only a portion of MG are reprogrammed into retinal neurons, whereas the rest are quiescent or maintain glial identity. In zebrafish, three MG populations expressing Stat3 and Ascl1a and possessing different regenerative capacities have been screened following retinal injury^29^. Another study defined two MG subpopulations based on the opposing role of Fgf8a on Notch signaling in zebrafish ^30^. scRNA-seq of MG in zebrafish revealed heterogeneity of MG and identified different reactive and non-reactive MG subpopulations ^31^. In avians, MG in the central and peripheral retinas maintain different regenerative capacities in response to damage, and MG subpopulation with elevated level of the glial fibrillary acidic protein (GFAP) cannot be reprogrammed into retinal progenitor cells^32^. In mammals, some progenitor-associated transcription factors, such as Chx10 and Pax6, are expressed only in a portion of MG^33^. However, whether MG in mammals consist of diverse MG subpopulations with different regenerative capacities remains unclear.

Here, we determined the changes in MG in RD 10 (rd10) mice from pre- to end-stage degeneration. One distinctive MG subpopulation (Chx10^−^Pax6^+^ MG) was activated following the initiation of degeneration and migrated to the outer nuclear layer (ONL), whereas Chx10^+^ MG and Pax6^−^ MG remained in the inner nuclear layer (INL). All of MG failed to reprogram into photoreceptors (PRs). Single-cell RNA sequencing (scRNA-seq) of MG revealed 9 subclusters, in which Chx10^+^ MG subclusters were related to synapse and regulation of neuron death and Chx10^−^ MG subclusters were related to cell migration, stem cell population and visual perception. hAECs transplantation induced extensive migration of the specific MG subpopulation (Chx10^−^Pax6^+^ MG) into the ONL from the INL and reprogrammed them into PR-like cells. Mechanistically, hAECs facilitated MG reprogramming through Mmp2- and Mmp9-mediated ECM remodeling possibly.

## Results

### Specific MG subpopulations are activated and migrate into ONL in degenerative retinas

We first investigated the changes in MG caused by the expression of GFAP and SRY-box 9 (Sox9) (a marker of MG nuclei) in C57 and rd10 mice. In the retinas of C57 mice, GFAP was expressed only in astrocytes at the ganglion cell layer (GCL) and was absent in MG (Figure S1a). In the retinas of rd10 mice at 2 weeks postnatal (2 wpn) before PR degeneration, MG were activated and expressed GFAP. With the progression of degeneration, GFAP levels increased, and glial scar formed at the outer limiting membrane at 8 wpn when degeneration progressed to the end stage (Figure S1a, b). The number of Sox9^+^ nuclei in the entire retina remained constant during the degeneration stage (Figure S1c). Migration of Sox9^+^ nuclei was first observed in the retinas of rd10 mice at 3 wpn, which was later than GFAP expression onset in MG at 2 wpn (Figure 1a). The number of Sox9^+^ nuclei that migrated into the ONL increased with degeneration, peaking at 6 and 8wpn (Figure 1b).

**Figure 1.**
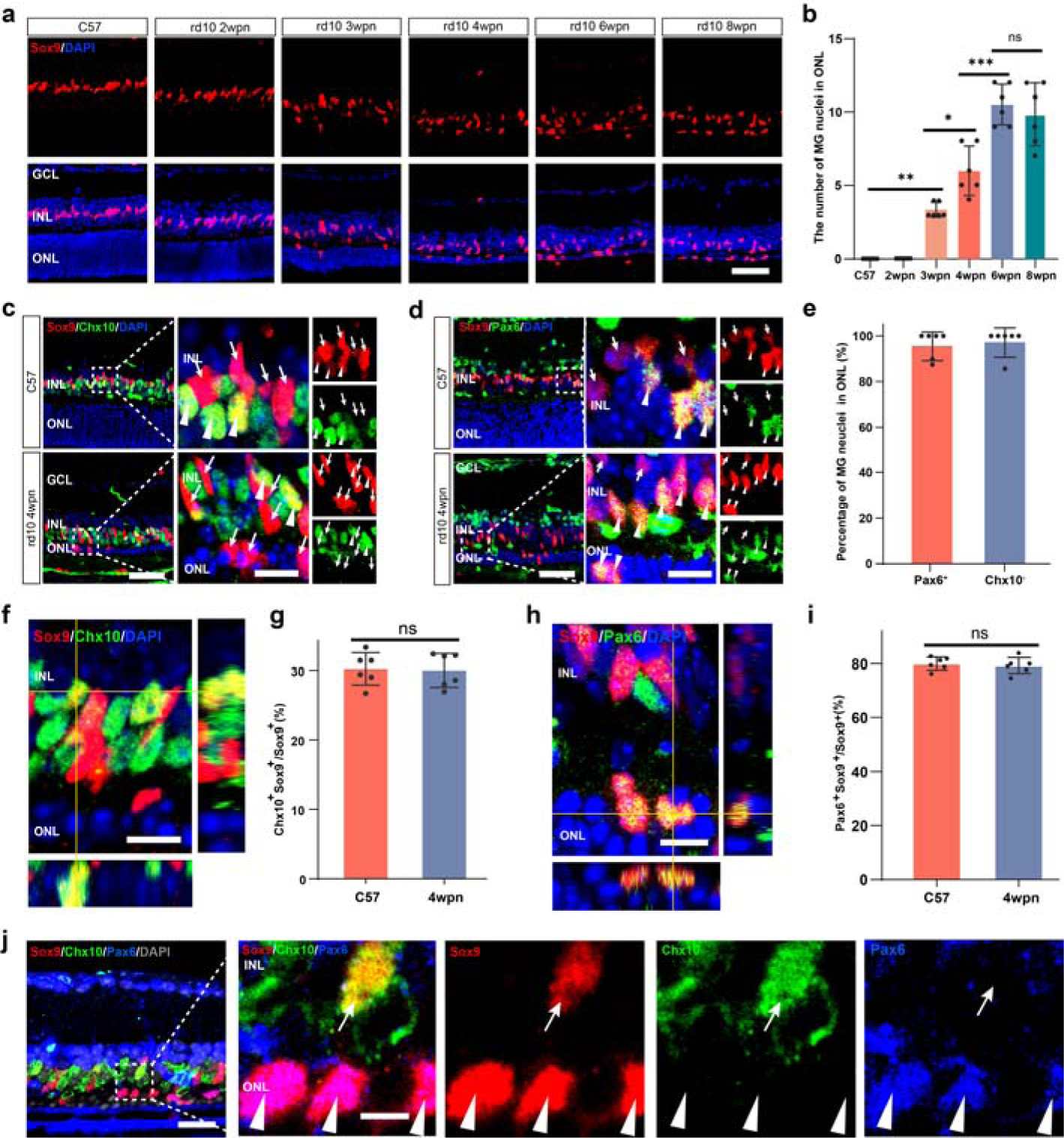
Müller glia (MG) subpopulation migrated to the ONL in the degenerative retina. **a, b** Representative immunofluorescence images and number of Sox9^+^ nuclei in the ONL of C57 and rd10 mice from 2 weeks postnatal (wpn) to 8 wpn. **c** Representative immunofluorescence images of Sox9 and Chx10 in the retinas of rd10 mice at 4 wpn, arrow: Sox9^+^Chx10^−^ nuclei, arrowhead: Sox9^+^Chx10^+^ nuclei. **d** Representative immunofluorescence images of Sox9 and Pax6 in the retinas of rd10 mice at 4 wpn, arrow: Sox9^+^Pax6^−^ nuclei, arrowhead: Sox9^+^Pax6^+^ nuclei. **e** Percentage of Chx10^−^ and Pax6^+^ in Sox9^+^ nuclei in ONL of rd10 mice. **f** Orthogonal image of Sox9 and Chx10 co-labeling in rd10 mice at 4 wpn. **g** Percentage of Sox9^+^ and Chx10^+^ nuclei in total Sox9^+^ nuclei at 4 wpn. **h** Orthogonal image of Sox9 and Pax6 co-labeling in rd10 mice at 4 wpn. **i** Percentage of Sox9^+^ and Pax6^+^ nuclei in total Sox9^+^ nuclei at 4 wpn. **j** Graph showing immunofluorescence staining of Sox9, Chx10, and Pax6. Arrow: Sox9^+^ and Chx10^+^ nuclei; Arrowhead: Sox9^+^ and Pax6^+^ nuclei. Data are presented as mean ± SD, n = 6. For statistical analysis, one-way ANOVA followed by Tukey’s multiple comparison tests (b) and Students’ *t* test (g, i) are applied. * *p* < 0.05, ** *p* < 0.01, *** *p* < 0.001; ns, no significance. Scale bar, 50 μm (a, c, d, j), 10 μm (enlarged images of c, d; f, h), 5 μm (enlarged images of j).

To investigate whether MG subpopulations exist, we co-stained Sox9 with two retinal progenitor markers, Chx10 and Pax6. In the retinas of C57 mice, Sox9^+^ nuclei consisted of Sox9^+^Chx10^+^ (arrowhead) and Sox9^+^Chx10^−^ nuclei (arrow) (Figure 1c). In degenerative retinas at 4, 6, and 8 wpn, Sox9^+^Chx10^−^ nuclei (arrow) migrated to the ONL, while Sox9^+^Chx10^+^ nuclei (arrowhead) remained in the INL (Figure 1c, Figure S1d). Correspondingly, Sox9^+^Pax6^+^ (arrowhead) and Sox9^+^Pax6^−^ nuclei (arrow) were located in the INL of C57 mice, whereas in rd10 mice, Sox9^+^Pax6^+^ nuclei (arrowhead) migrated to the ONL, and Sox9^+^Pax6^−^ nuclei (arrow) remained in the INL (Figure 1d). Sox9^+^ nuclei that migrated into ONL were almost Pax6^+^ and Chx10^−^ (Figure 1e). Orthogonal views confirmed the co-labeling of Sox9 with Chx10 and Pax6 (Figure 1f, h). However, the percentages of Sox9^+^Chx10^+^ or Sox9^+^Pax6^+^ nuclei in the total Sox9^+^ nuclei in C57 and rd10 mice were almost equal (Figure 1g, i). In addition, multiplex immunofluorescence images demonstrated that Sox9^+^Chx10^−^Pax6^+^ nuclei (arrowhead) migrated to the ONL, while Chx10^+^ and Pax6^−^ nuclei remained in the INL (Figure 1j). Given that Chx10 and Pax6 are expressed in of bipolar and amacrine cells, respectively, we co-labeled Sox9 with PKCa and HuC/D and found that all Sox9^+^ cells were MG and not neurons (Figure S1f, g). We investigated the final fate of MG that migrated to the ONL by staining with rhodopsin (a PR marker). Sox9 staining was not co-labeled with rhodopsin staining in the ONL (Figure S1e). Thus, we confirm that distinctive Chx10^−^Pax6^+^ MG subpopulation exist in healthy retinas. During RD, the Chx10^−^Pax6^+^ MG subpopulation is activated and migrates to the ONL, whereas the other subpopulations remain in the INL.

### scRNA-seq analysis of MG in healthy and degenerative retinas

To further study the molecular and functional signatures of MG subpopulations, we isolated tdTomato^+^ MG from MG lineage-tracing mice at 8 wpn for scRNA-seq. We integrated our data with published scRNA-seq data obtained from healthy retinas and rd10 mice at 3 wpn using Harmony after removing unqualified cells through quality control ^34–36^ (Figure S2a-c). Unsupervised clustering revealed the main cell types with a unique gene expression profile in the retina and identified various cell types (Figure S2d-f). We analyzed the differences in gene expression patterns of cells between adult retina and developmental retina ^37^ and and observed that among the cell types in adult retina, cell similarity between MG and retinal progenitor cells (early retinal progenitors, late retinal progenitors and neurogenic cells) in developmental retinal was the highest (Figure S2g). According to the gene set based on the gene expression profiles of retinal progenitor cells (RPCs)^37^(Supplemental table 1), the RPC score of MG was the highest among all cell types in the retina (Figure S2h). After removing other cell types, 12600 MG from healthy and degenerative retinas remained (Figure S2i). Re-clustering identified 9 MG subclusters (M1– M9), as shown by the Uniform Manifold Approximation and Projection (UMAP) plot (Figure 2a, Figure S2k, i). We used ROGUE to assess the purity of single cell clusters, the ROGUE scores indicated that cell composition of MG showed heterogeneity (Figure 2b) and MG subclusters showed higher purity (Figure 2c). We determined classifiers of MG subclusters using ROC analysis to classify different MG subclusters and AUC values (>0.8) of classifying genes for each subcluster indicated the differences between MG subclusters (Figure S2j). Violin plot showed the marker genes of each MG subcluster (Figure 2d). We found that Pax6 was expressed in every MG subclusters and Chx10 was highly expressed only in M5 and M6 subclusters (Figure 2e). Multiple comparisons revealed that Chx10 expressions in M5 and M6 were higher than those in other subclusters significantly (Figure 2f). Thus, M5 and M6 were defined as Chx10^+^ subclusters and other subclusters were Chx10^−^. GSEA showed that Chx10^+^ subclusters (M5 and M6) upregulated the activity of modulation of chemical synaptic transmission and regulation of neuron death (Figure 2g, h). Gene Ontology (GO) enrichment analysis of the marker genes of each MG subclusters was used to explore their potential functions (Figure S3a). Among Chx10^−^MG subclusters, M1 was related to cell migration, gliogenesis, neurogenesis, stem cell maintenance, and wnt signaling (Figure 2i). M3 and M4 were related to visual perception, light stimuli ATP metabolism and electron transport chain, (Figure 2j, k). GSEA further confirmed that the activities of above-mentioned pathways were upregulated in M1, M3 and M4 respectively (Figure S3b-h). SCENIC analysis showed that transcription factors related to neuron development and photoreceptor development including Crx, Etv1, Gatad1 were activated in M1, M3 and M4 (Figure S3i, j). Besides, M5 and M7 might play an important role in regulation of retinal immune response and neuron death during retinal degeneration (Figure S3a).

**Figure 2.**
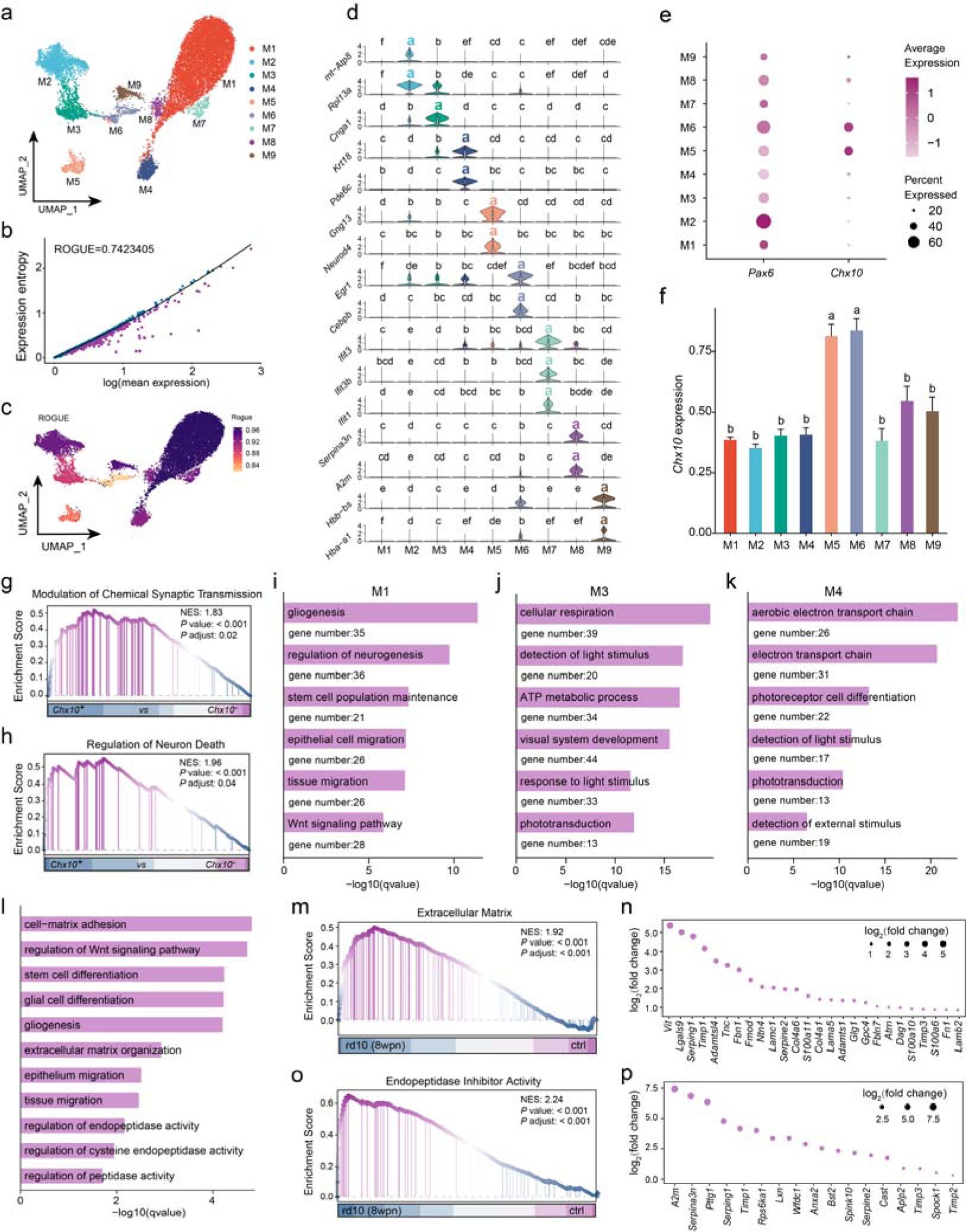
scRNA-seq analysis for diverse Müller glia (MG) subtype and differential analysis between MG from healthy and degenerative retinas. **a** UMAP plot showing the unsupervised subclustering of MG from healthy and degenerative retinas colored by subcluster. **b, c** ROGUE analysis showing the cell purity of MG and MG subclusters. **d** Violin plot showing the expressions of markers for each MG subclusters. **e** Dotplot showing expressions of Pax6 and Chx10 in different MG subclusters. **f** Diagram showing the expressions of Chx10 in M5 and M6 is significantly higher than others. Data are presented as mean ± SEM. **g, h** GSEA of Chx10^+^ (M5, M6) and Chx10^−^ MG subclusters. **i-k** GO enrichment analysis of marker genes of M1, M3 and M4. **l** GO analysis of DEGs between healthy and degenerative retinas in MG. **m, o** GSEA showing extracellular matrix (m) and endopeptidase inhibitor activity (o) gene sets in MG from healthy and degenerative retinas. **n, p** Point plot showing expression changes of genes in extracellular matrix (n) and endopeptidase inhibitor activity (p) in degenerative MG compared to healthy MG. Point size indicates log2(fold change). For statistical analysis of d and f, one-way ANOVA followed by Tukey’s multiple tests is applied and plots marked with the different letter have statistically significant differences (*p* value < 0.05).

Differential expression analysis between MG from healthy and degenerative retinas at 3 wpn showed that differentially expressed genes (DEGs) were related to gliogenesis, cell migration, interferon signaling and peptidase activity (Figure S4a, b). GSEA confirmed that the activities of cell migration, peptidase inhibitory activity, immune response, and interferon signaling were upregulated in MG in degenerative retinas (Figure S4c-h). At 8 wpn, GO terms of DEGs between MG from healthy and degenerative retinas included gliogenesis, cell migration, immune responses, neuron death and wnt signaling (Figure 2l, Figure S4i, j). Terms related to gliogenesis, regulation of chromatin organization, regulation of glial cell migration were upregulated and terms related to respiratory chain complex, ribosome were downregulated in MG from degenerative retinas (Figure S4k, l, n-p). More importantly, using GSEA, we found that extracellular matrix, integrin-mediated signaling pathway and endopeptidase inhibitor activity were upregulated in MG from degenerative retinas (Figure 2m, o, Figure S4m). ECM genes including Vit, Tnc, Col4a6, Lamc1, Lama5, Glg1, and peptidase inhibitor activity genes including A2m, Serpina3n, Timp1, Timp2 were upregulated in MG from degenerative retinas (Fig 2n, p).

Therefore, the scRNA-seq data confirm the heterogeneity of MG in mammal retinas and reveal that Chx10^+^ MG subclusters were related to synaptic transmission and neuron death, Chx10^−^ MG subclusters were related to cell migration, stem cell maintenance and visual perception. Differential analysis suggests that MG in the degenerative retinas upregulate pathways of regulating ECM remodeling.

### hAECs promote MG subpopulation migration into ONL

To investigate whether hAECs could induce specific MG subpopulation reprogramming, we transplanted them into the subretinal space of rd10 mice at 2 wpn and administered phosphate-buffered saline (PBS) as the control (Figure S5a). The hAECs isolated from the placenta presented typical cobblestone morphology under a light microscope (Figure S5b). For tracing transplanted cells, hAECs were labeled with enhanced green fluorescent protein (EGFP) using cytomegalovirus promoter-driven EGFP lentivirus, resulting in more than 80% of hAECs were being labeled (Figure S5c, d). hAECs aggregated at the injection site after transplantation and were widely distributed throughout the retinal areas 1-week post-transplantation (wpt) (Figure S5e, f). Optical coherence tomography imaging showed that the transplanted hAECs were distributed in the subretinal space of rd10 mice at 1 wpt (Figure S5g).

The number of Sox9^+^MG nuclei in the entire retina was higher in the hAECs than in the PBS group, implying that hAECs might promote MG proliferation (Figure 3a, b). There was no statistically significant difference in the total number of MG in the entire retina in the hAECs group at 2, 4, and 6 wpt (Figure 3c). In the retinas of rd10 mice at 2 wpt, only a few Sox9^+^ nuclei migrated to the ONL and were located on the inner side of the ONL (Figure 3a, d). In the retinas of age-matched transplanted mice, large numbers of Sox9^+^ nuclei migrated into the ONL (Figure 3a, d). In the retinas of mice at 4 wpt, Sox9^+^ nuclei migration to the ONL was higher in the hAECs group than that in the PBS group; similar changes were observed in the retinas at 6 wpt (Figure 3a, d). The number of Sox9^+^ nuclei that migrated to the ONL constantly increased from 2 to 4 wpt, but this increase was not statistically significant at 6 wpt compared with that at 4 wpt (Figure 3e). The Sox9^+^ nuclei migration distance (the distance from the midline of the INL to the Sox9□ nuclei located in the ONL) also increased over time following hAECs transplantation (Figure 3f, g). In the hAECs group at 2 wpt, most of the migrated Sox9^+^ nuclei were on the inner side of the ONL, while several were on the lateral side (Figure 3a). Sox9^+^ nuclei migrated to the middle of the ONL at 4 wpt, and were distributed throughout the ONL at 6 wpt. Similarly, the MG subpopulation stained with Chx10 and Pax6 showed that the Sox9^+^ nuclei migrated to the ONL were Chx10 negative and Pax6 positive, which was further confirmed by orthogonal views (Figure 3h-k).

**Figure 3.**
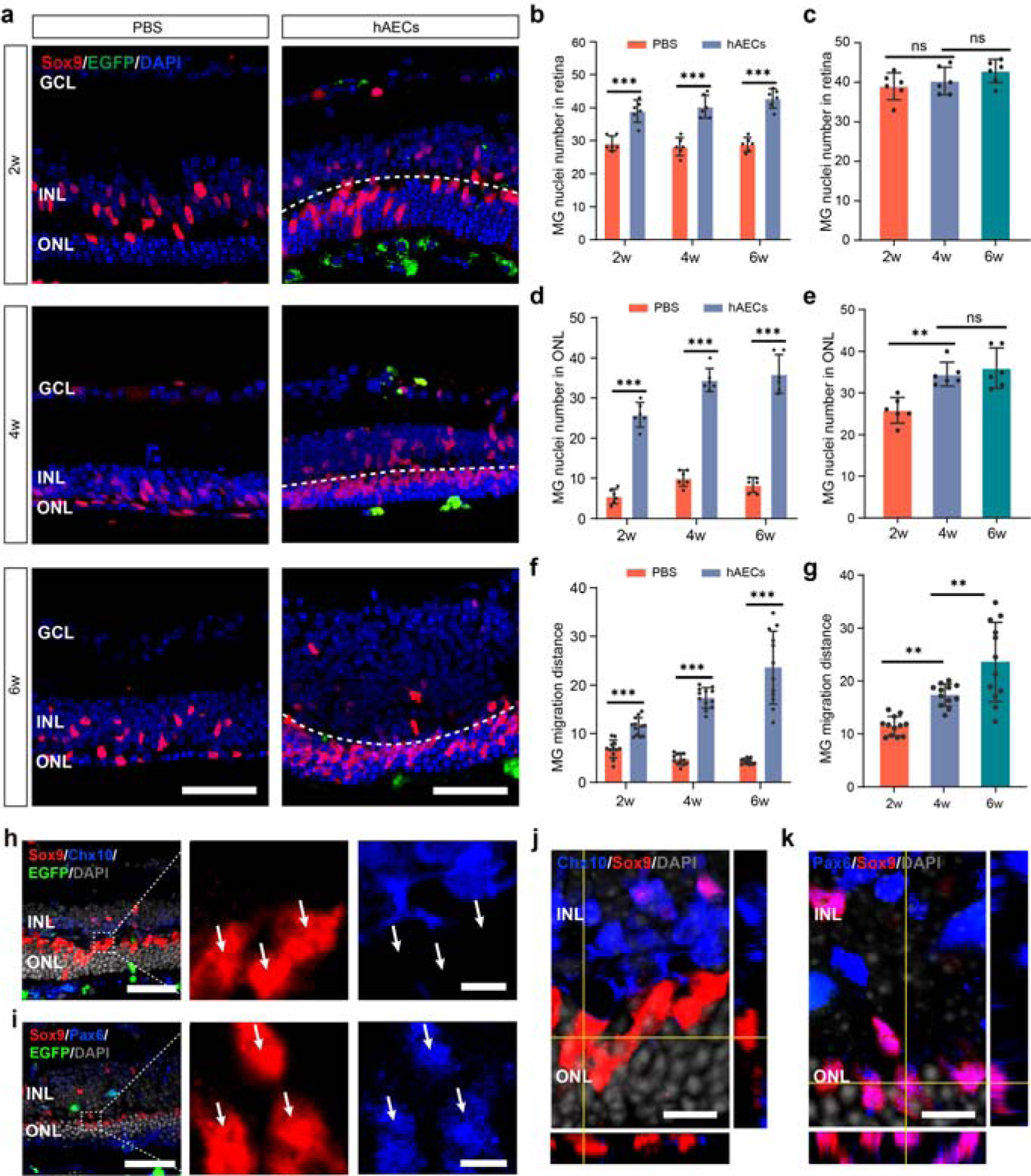
Transplanted hAECs promote Müller glia (MG) subtype migration to the outer nuclear layer (ONL). **a** Representative immunofluorescence images of Sox9 at 2, 4, and 6 weeks post-transplantation (wpt) in the PBS and hAECs groups. **b** Quantitative analysis of the number of Sox9^+^ nuclei in the whole retina. **c** Quantitative analysis of the number of Sox9^+^ nuclei in the whole retina in the hAECs group. **d** Quantitative analysis of the number of Sox9^+^ nuclei that migrated to the ONL. **e** Quantitative analysis of the number of Sox9^+^ nuclei that migrated to the ONL in the hAECs group. **f** Quantitative analysis of MG migration distance. **g** Quantitative analysis of MG migration distance in the hAECs group. **h** Representative immunofluorescence images of Sox9 and Chx10 in the hAECs group at 2wpt. **i** Representative immunofluorescence images of Sox9 and Pax6 in hAECs group at 2wpt. **j** Orthogonal views of Sox9 (red) and Chx10 (blue) staining. **k** Orthogonal views of Sox9 (red) and Pax6 (blue) staining. Data are presented as mean ± SD, n = 6. For statistical analysis, one-way ANOVA followed by Tukey’s multiple tests (c, e, g) and multiple *t* test with Benjamini-Hochberg correction (b, d, f) are applied. * *p* < 0.05, ** *p* < 0.01, *** *p* < 0.001; ns, no significance. Scale bars, 50 μm (a, h, i), 10 μm (j, k), 5 μm (enlarged images of h, i).

To verify whether hAECs promote MG proliferation and directional migration, we performed *in vitro* experiments using the human MG cell line MIO-M1 (figure S6a, b). MIO-M1 cells were Sox9^+^/Pax6^+^/Chx10^−^, which correspond to MG subpopulation that migrated into ONL in the degenerative retinas (Figure S6a). MIO-M1 cells were characterized by a flattened morphology with broad lamellipodia under normal conditions (Figure S7a). MIO-M1 cells co-cultured with hAECs underwent a morphological shift, showing a reduced cell area and an increased cell process length and elongation index (Figure S7a-d)^38^. The cell cycle test showed that the percentage of cells in the S phase increased in the hAECs group, while the cell migration assay showed that the number of migrated cells increased in the hAECs group (Figure S7e-j).

These results indicate that hAECs promote MG proliferation and trigger Sox9^+^/ Pax6^+^/Chx10^−^ MG migration to the ONL.

### hAECs induce migrated MG reprogramming into photoreceptor-like cells

Next, we investigated the final fate of the MG that migrated to the ONL. Compared with the C57 group, the expression of GFAP in degenerative retinas increased, and a glial scar in the outer retina was formed (Figure 4a, b). Notably, GFAP expression was reduced and almost disappeared from the outer retina after hAECs transplantation (Figure 4a, b), which was also confirmed using western blotting (Figure 4c, d). Furthermore, we observed co-labeling of Sox9 and rhodopsin in the hAECs group (Figure 4e). DAPI staining of the C57 mice retinas revealed that the MG and PRs exhibited distinct nuclear architectures^39,40^ (Figure S8a). The conventional nuclear pattern of MG was observed in the degenerative retina (Figure S8b). Notably, some of the Sox9^+^ nuclei that migrated to the ONL in the hAECs group transitioned into PR-like nuclei, which was verified using the orthogonal view, which was in line with the nuclear architecture changing as RPCs differentiated into photoreceptors during development^39^ (Figure 4f-h). The strongly DAPI-positive central chromocenter was stained pale for Sox9, suggesting that Sox9 protein might be excluded from the nuclei, and MG were losing glial identity (Figure 4f).

**Figure 4.**
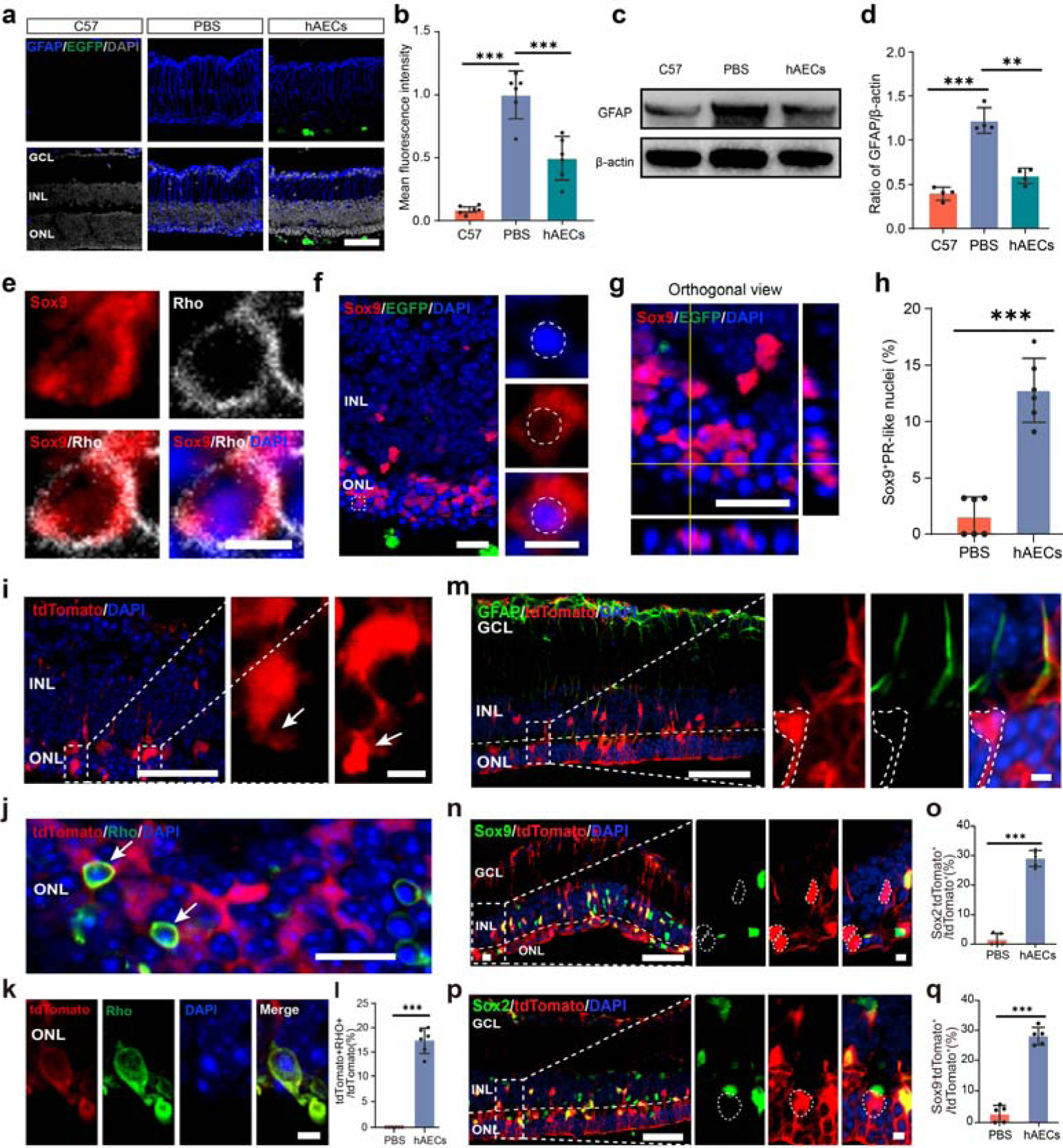
Transplanted hAECs induced Müller glia (MG) to lose glial markers and reprogram them into photoreceptors. **a, b** Representative immunofluorescence images and relative quantitative analysis of GFAP at 6 weeks post-transplantation (wpt). **c, d** Western blotting results and quantitative analysis of GFAP at 6 wpt. **e** Representative immunofluorescence images of Sox9 and rhodopsin co-labeling in the hAECs group. **f, g** Representative images and orthogonal views of Sox9 staining and nuclear architecture at 6 wpt in the hAECs groups. **h** Percent of Sox9^+^ and photoreceptor-like nuclei in total Sox9D nuclei in the PBS and hAECs groups. **i** Representative images of tdTomato^+^ MG morphology that migrated into ONL in MG-tracing mice. Arrow indicated the outer segment-like structure. **j** Representative images of Rhodopsin and tdTomato in hAECs group at 6 wpt. **k** Co-labeling of Rhodopsin and tdTomato in MG with photoreceptor-like morphology. **l** Quantitative analysis of Rhodopsin^+^/tdTomato^+^ MG in the outer nuclear layer at 6 wpt. **m** Representative images of tdTomato and GFAP staining at 6 wpt. **n-q** Representative images and percentage of Sox9^−^ (**n, o**) and Sox2^−^ **(p, q**) nuclei in all tdTomato^+^ nuclei at 6 wpt. Data are presented as mean ± SD, n = 6, n = 5 (o, q), n = 4 (d). For statistical analysis, one-way ANOVA followed by Tukey’s multiple tests (b, d) and Students’ *t* test (h, i, o, q) are applied. * *p* < 0.05, ** *p* < 0.01, *** *p* < 0.001; ns, no significance. Scale bars, 50 μm (a, i, m, n, p), 20 μm (f, g), 5 μm (e, k, enlarged images of f, i, m, n, p).

MG lineage-tracing mice were used to confirm the reprogramming of MG into PR-like cells. In the retinas of MG-lineage-tracing mice, approximately all tdTomato^+^ cells were Sox9^+^, indicating that tdTomato accurately labeled MG (Figure S8c). In the hAECs group, several MG that migrated into ONL generated photoreceptor outer segment (POS)-like structure (Figure 4i). MG in the ONL were partly positive for rhodopsin and MG with a POS-like structure were rhodopsin^+^, whereas no MG expressed rhodopsin in the PBS group (Figure 4j-l). Meanwhile, tdTomato^+^ MG that migrated to the ONL were negative for glial markers, including GFAP, Sox9, and Sox2, in the hAECs group, whereas almost no MG lost glial markers in the PBS group (Figure 4m-q). Similarly, the nuclear architecture of MG following hAECs transplantation displayed a PR-like nuclear pattern; this was not the case in the degenerative retina (Figure S8d-f). Based on the morphological features of the nuclear architecture, the progression of PR-like nuclei derived from MG was divided into four stages: rest, activated, intermediate, and terminal (Figure S8f, g). In addition, MG in the ONL were not co-labeled with other neuronal markers, including HuC/D (amacrine and retinal ganglion cells) and PKCa (bipolar cells), suggesting they did not differentiate into neurons other than PRs (Figure S8h, i). Considering that previous research has reported MG-engulfing PRs, we examined whether MG reprogramming into PR-like cells was caused by phagocytosis using the lysosomal marker Lamp1. The results showed that MG that migrated into the ONL were hardly co-labeled with Lamp1 at 6wpt (Figure S8j, k)^41^. Therefore, we conclude that MG in the ONL are reprogrammed into PR-like cells rather than other retinal neurons following hAECs transplantation.

### hAECs potentially regulate ECM remodeling by regulating Mmp9 and Mmp2

Excessive deposition of ECM components such as chondroitin sulfate proteoglycans (CSPGs) is frequently associated with CNS damage, including RD, which is a critical barrier to neuronal repair and regeneration^42–45^. On the one hand, GO and Kyoto Encyclopedia of Genes and Genomes (KEGG) analysis of the top 200 genes in hAECs showed that pathways related to ECM regulation including regulation of cell-substrate adhesion, collagen metabolic process, regulation of peptidase activity and ECM-receptor interaction were enriched (Figure S9a, b)^46^; on the other hand, hAECs with high Mmp2 and Mmp9 expression degrade ECM components during parturition^47^. Thus, we investigated whether transplanted hAECs degrade inhibitory ECM components via secreting Mmp2 and Mmp9 to induce retinal regeneration. Consistent with our findings, ECM components, including CSPGs and collagen Ι, were deposited in the degenerative retina (Figure 5a-d)^42^. Almost no CSPGs were found in the retinas of C57 mice, whereas a clear CSPG band was deposited in the ONL and the external limiting membrane (ELM) (Figure 5a, b). Following hAECs transplantation, CSPGs deposited in the ONL and ELM were degraded (Figure 5a, b). Excessive collagen I deposition in the degenerative retina was reversed following hAECs transplantation (Figure 5c, d). To elucidate the mechanism by which hAECs regulate ECM remodeling, we investigated the expression levels of Mmp9 and Mmp2 in the retinas of the C57, PBS, and hAECs groups. Mmp9 was mainly expressed in the GCL and INL of the retinas of the C57 and PBS groups (Figure 5e). In the retinas of the hAECs group, increased Mmp9 expression was observed in OPL and ELM, where ECM components, including CSPGs and collagen I, were present (Figure 5e, g). The expression pattern of Mmp2 in the retinas of the C57, PBS, and hAECs groups was similar to that of Mmp9 (Figure 5f, h), and western blotting confirmed increased Mmp2 expression in the hAECs group relative to that in the other groups (Figure 5i, j). Therefore, we demonstrate that hAECs degrade inhibitory ECM components, including CSPGs and collagen Ι, via regulating Mmp9 and Mmp2, which might create a pro-regeneration microenvironment to induce MG reprogramming probably.

**Figure 5.**
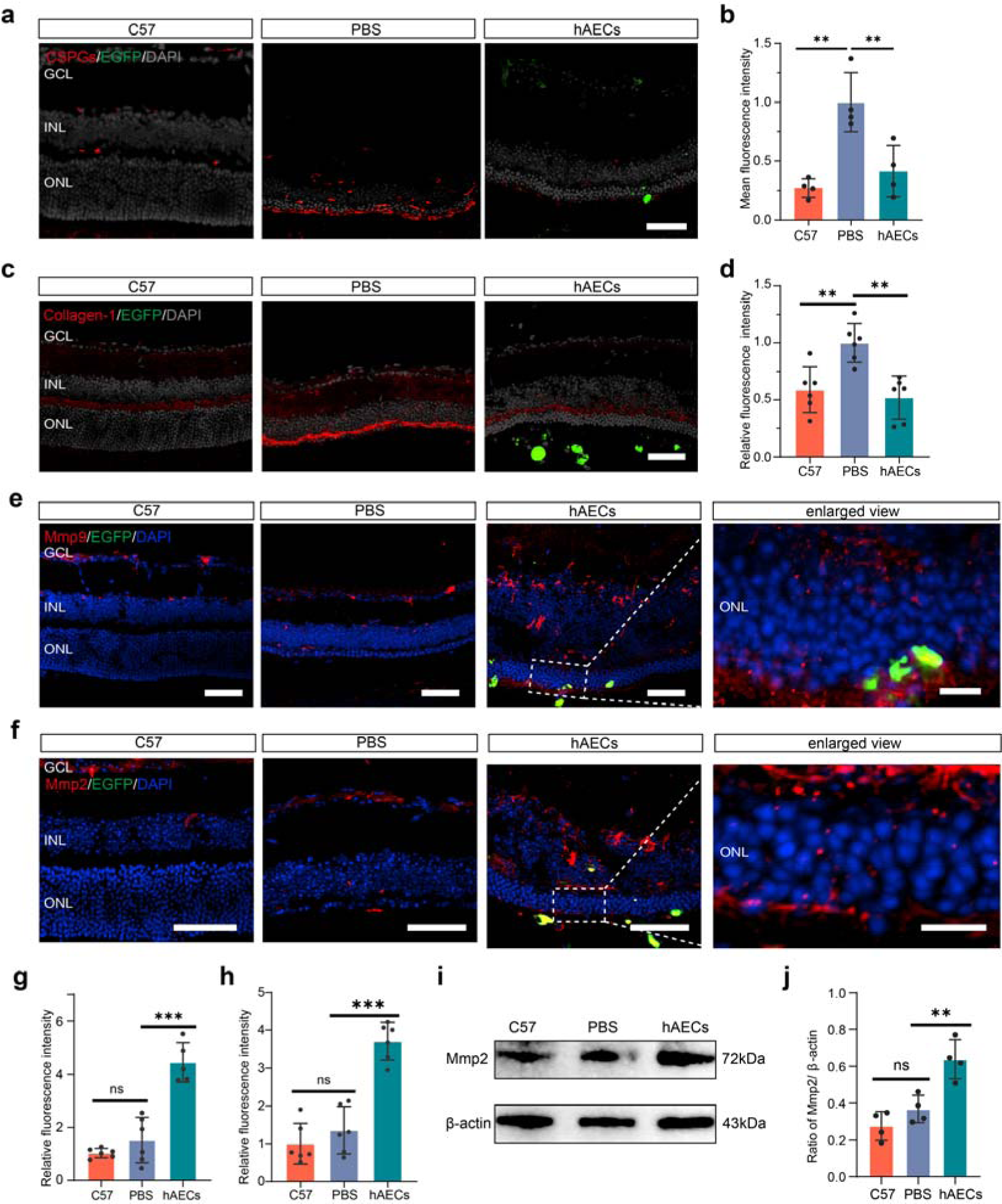
hAECs regulate extracellular matrix remodeling through secreted Mmp2/Mmp9 signaling. **a, b** Representative fluorescence images and relative quantitative analysis of chondroitin sulfate proteoglycans in the C57, PBS and hAECs groups at 6 weeks post-transplantation (wpt). **c, d** Representative fluorescence images and relative quantitative analysis of Collagen in the C57, PBS and hAECs groups at 6 wpt. **e, f** Representative fluorescence images of Mmp9 (E) and Mmp2 (F) in the C57, PBS and hAECs groups at 6 wpt. **g, h** Relative quantitative analysis of Mmp9 (G) and Mmp2 (H) in the C57, PBS and hAECs groups at 6 wpt. **i, j** Western blotting results of Mmp2 protein levels in retinas from the C57, PBS and hAECs groups. Data are presented as mean ± SD, n = 6 (d, g, h), n = 4 (b, j). For statistical analysis, one-way ANOVA followed by Tukey’s multiple tests (b, d, g, h, j) is applied. * *p* < 0.05, ** *p* < 0.01, *** *p* < 0.001; ns, no significance. Scale bars, 50 μm (a, c, e, f), 10 μm (enlarged images of e, f).

### hAECs protect retinal structure and function in rd10 mice

Finally, we investigated whether retinal structure and function in rd10 mice were rescued after hAECs transplantation. Compared with that in C57 mice, the expression of two PR markers, rhodopsin and cone-arrestin, in the PBS group drastically declined but was partially reversed in the hAEC group at 2, 4, and 6 wpt (Figure 6a). Analysis of ONL thickness also indicated that transplanted hAECs significantly protected PRs from degeneration at 2, 4, and 6 wpt (Figure 6b-d). Subsequently, electroretinography (ERG) was used to determine visual function in the C57, PBS, and hAECs groups and showed that the *b*-wave amplitudes in the hAECs group at different light intensity stimuli increased significantly compared with those in the PBS groups at 2, 4, and 6 wpt (Figure 6e-h). Moreover, hAECs rescued *a*-wave amplitude at multiple intensities at 2, 4, and 6 wpt (Figure 6i-k). In addition, the light/dark transition test showed that hAECs transplantation improved the visual behavior of the degenerative mice (Figure S10a-d). Thus, hAECs transplantation preserves retinal structure and improves visual function in rd10 mice.

**Figure 6.**
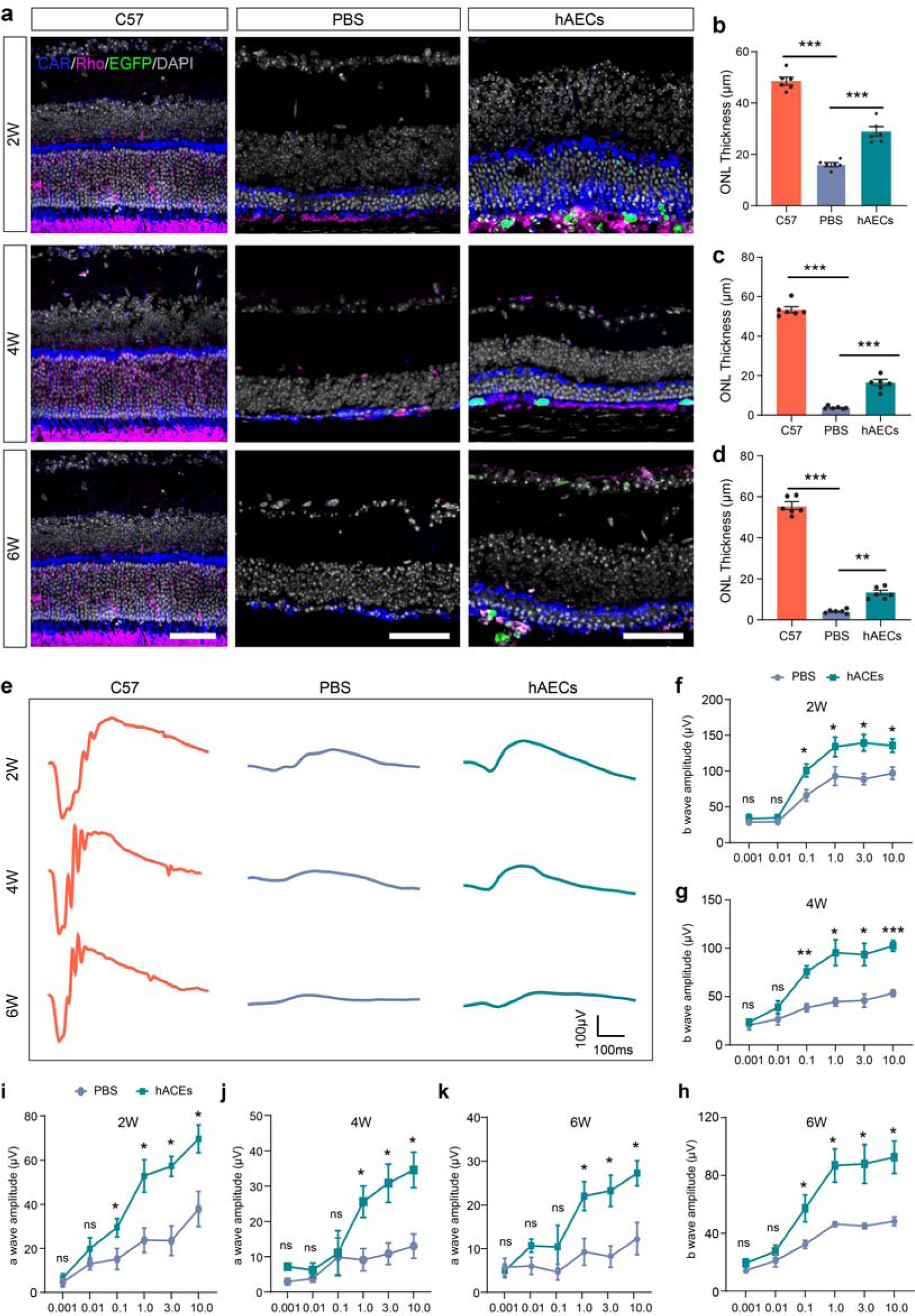
Transplanted hAECs protect photoreceptors and improve visual function in rd10 mice. **a** Representative immunofluorescence images of rhodopsin and cone-arrestin (CAR) at 2, 4, and 6 weeks post-transplantation (wpt). **b-d** Statistical analysis of outer nuclear layer (ONL) thickness at 2, 4, and 6 wpt. **e** Representative waveforms of the C57, PBS and hAECs groups at 2, 4, and 6 wpt through scotopic ERG at 3.0 log(cd*s/m^2^). **f-h** Statistical analysis of ERG *b*-wave amplitude at 2, 4, and 6 wpt at different light intensity stimuli. **i-k** Statistical analysis of ERG *a*-wave amplitude at 2, 4, and 6 wpt at different light intensity stimuli. Data are presented as mean ± SEM, n = 6 (b, c, d, f, i), n = 5 (g, j, h, k). For statistical analysis, one-way ANOVA (b, c, d) followed by Tukey’s multiple tests and multiple *t* test with Benjamini-Hochberg correction (f-k) are applied. * *p* < 0.05, ** *p* < 0.01, *** *p* < 0.001; ns, no significance. Scale bars, 50 μm.

## Discussion

Reprogramming MG to regenerate retinal neurons is a promising treatment for RD, which is characterized by retinal neuron loss. Determining MG heterogeneity is critical for accurately defining its functions and utilizing MG reprogramming to regenerate retinal neurons. In the present study, we investigated MG heterogeneity in healthy and degenerative retinas on a histological and single-cell basis. We found that hAECs transplanted into rd10 mice induced a specific MG subpopulation to migrate and reprogram into PR-like cells by regulating ECM remodeling.

We defined a distinctive MG subpopulation in the retinas of mammals: Sox9^+^Chx10^−^Pax6^+^ MG. During RD, a few Chx10^−^Pax6^+^ MG were activated and migrated to the ONL, whereas Chx10^+^ MG and Pax6^−^ MG remained in the INL. Following hAECs transplantation, most of the Chx10^−^ MG subpopulation migrated to the ONL and were reprogrammed into PRs, indicating that Chx10 and Pax6 have different roles in determining MG fate. In the adult retina, Chx10 is primarily expressed in bipolar cells, whereas Pax6 is expressed in amacrine and retinal ganglion cells^9^. However, these two transcription factors play different roles in maintaining the pluripotency of RPCs and regulating neuronal differentiation during retinal development^48,49^. Pax6 is expressed in all RPCs and is recognized as the most critical transcription factor during eye development, especially retinal development^49^. Pax6 has been detected in the eye field at embryonic day 8, is strongly expressed in all cells of the optic cups, and is responsible for regulating other transcription factors, including Atoh7, Mash1, Neurod1, Ngn2, and Crx^48,50^. Moreover, Pax6 in RPCs is essential for maintaining their pluripotency and proliferation, and its deficiency leads to failure of neuronal differentiation in retinal ganglion cells, PRs, bipolar cells, and horizontal cells^51,52^. Notably, Pax6 overexpression converted human embryonic stem cells (hESCs) into neural lineage cells^53^. Although Chx10 is also a marker of RPCs, it is expressed later than Pax6 and is first expressed in RPCs with low Pax6 expression^48^. Previous research has reported that Chx10 overexpression increases bipolar cells but decreases PR differentiation, whereas Chx10 knockdown has the opposite effect^54^. These results imply that Chx10 regulates RPC differentiation into bipolar cells rather than PR cells, which may explain why only the Chx10^−^ MG subpopulation was reprogrammed into PR-like cells following hAECs transplantation.

scRNA-seq showed that the MG contained Chx10^+^ and Chx10^−^ subclusters. Chx10^+^ MG subclusters M5 and M6 were related to synaptic transmission and regulation of neuron death, which might be MG that remained in INL during retinal degeneration. Among Chx10^−^ subclusters, M1 M3, and M4 subclusters were related to cell migration, and stem cell maintenance, visual perception and might be MG subpopulations that were reprogrammed into PR-like cells in the hAECs group. Notably, the M7 subpopulation highly expressed IFN-related genes, including Ifit1 and Ifit3. IFN from MG showed increased expression in the optic nerve crush injury model, peaking at injury midpoint, and played a vital role in coordinating the immune response after injury^55^. Thus, the M7 subpopulation may regulate immune responses in degenerative retinas^56^. Differential expression analysis in MG suggested that pathways related to cell migration, immune response were upregulated in degenerative retina. In addition, pathways and genes related to ECM were upregulated in MG from degenerative retina, which might imply the fibrosis phenotype of MG and suppress neural regeneration ^57^.

In the retinas of the hAECs group at 6 wpt, not all PRs were tdTomato^+^, indicating that a substantial number of original PR cells were shielded from death following hAECs transplantation. hAECs synthesize and release numerous neurotrophic and anti-inflammatory factors, including brain-derived neurotrophic factor, neurotrophin-3, nerve growth factor, and insulin-like growth factor, to protect neurons from various CNS diseases^58,59^. Therefore, we speculated that hAECs promoted MG reprogramming and protected PR cells during RD. We observed that the positive signal of Sox9 was transferred from the nucleus to the cytoplasm in the hAECs group and was gradually downregulated, which was also the case for Sox2. When qNSCs are activated to initiate neuronal fate specification, the expressions of Sox9 and Sox2 are downregulated^60^, which is consistent with our results. In addition, we speculated that MG engulfed PRs since they can engulf apoptotic PRs and degrade them in lysosomes during retinal development and degeneration^41,61^. However, the lysosome marker Lamp1 did not co-label tdTomato^+^ cells with a PR-like nucleus, suggesting that the reprogramming we observed was not caused by phagocytosis.

hAECs secrete matrix metalloproteinases that degrade the ECM to facilitate delivery^47^. Based on this characteristic, hAECs transplantation effectively treats fibrotic diseases, such as intrauterine adhesions and lung fibrosis. hAECs promote tissue repair and regeneration by regulating ECM remodeling via matrix metalloproteinase activity^26,62^. After CNS damage, inhibitory ECM components, such as CSPGs, are excessively deposited around the injured sites and hinder neural regeneration^45,63^. It is well documented that CSPG degradation via the regulation of Mmp2 activity promotes oligodendrocyte progenitor cell migration, maturation, and remyelination in multiple sclerosis models^64^. In spinal cord injury, Mmp2 suppressed CSPG deposition and glial scar formation, and facilitated astrocyte migration and functional recovery^65,66^. Consistent with this study, degradation of inhibitory ECM components, such as CSPGs, activated RhoA/Actin/YAP signaling in astrocytes, promoting neural regeneration following CNS injury^43^. Similarly, hAECs transplantation enhanced the activities of Mmp2 and Mmp9 and degraded inhibitory ECM components, including CSPGs, which may promote retinal regeneration from MG through RhoA and cytoskeleton signaling. Another ECM component, Collagen-Ι expression was reduced after hAECs transplantation. Upregulation of Collagen-Ι expression will increase the tissue stiffness and reduce tissue elasticity^67,68^. Thus, reduction in Collagen-Ι expression caused degenerative retina to become soft, which may facilitate MG reprogramming^69^. However, other ECM components, such as lamin-B1, can promote neurogenesis. The decline in lamin-B1 levels impairs NSC activity and accelerates astrocyte aging in the hippocampus, which impedes adult neurogenesis. As expected, lamin-B1 overexpression promotes neurogenesis in the aged hippocampus^70,71^.

In addition to regulating the remodeling of the extracellular matrix, hAECs could release a very large number of neurotrophic factors, growth factors, anti-inflammatory factors, and exosomes ^72,73^, which have been widely used to promote the cell survival in lung injury^74^, kidney injury^75^, liver disease^76^, and ocular surface diseases^77^. In brain injuries, hAECs promoted the survival of neurons and neurogenesis through paracrine effects^78^. Besides, hAECs played an important role in regulating the immune microenvironment and transplanted of hAECs significantly rescued the neuronal death through immunomodulation^79–81^. It is noteworthy that regulation of microglia-mediated neuroinflammation and other inflammatory cytokines also facilitated MG reprogram into retinal progenitors in mice^42^. It revealed that the other potential mechanism of hAECs besides ECM remodeling promoted MG reprogramming.

However, several key questions remain unanswered. The results of scRNA-seq have not yet been verified using fluorescence in situ hybridization and immunofluorescence staining. Although we observed the MG heterogeneity and MG reprogramming through hAEC transplantation in rd10 mice, more models of retinal pigment degeneration disease should be taken into consideration to confirm this conclusion. The functions and features of PR-like cells derived from MG need to be confirmed using patch-clamp techniques and further scRNA-seq. Whether direct ECM regulation induces MG reprogramming and the specific mechanisms regulating ECM-induced MG reprogramming, require further investigation.

## Methods

### Animals

C57BL/6J and rd10 mice were acquired from the Experimental Animal Center of Third Military Medical University (Army Medical University). Glast-CreERT C57BL/6J mice were purchased from the Jackson Laboratory (cat # 012586; Bar Habor, ME, USA) and crossed with H11-tdTomato C57BL/6J mice (H11^em1Cin(CAG-LoxP-ZsGreen-Stop-LoxP -tdTomato)^) to generate Glast-CreERT/H11-tdTomato mice. We then generated MG lineage-tracing mice with a RD phenotype by crossing Glast-CreERT/H11-tdTomato mice with rd10 mice^42^. All mice were raised in a specific pathogen-free room at the Animal Care Center of Southwest Hospital on a 12-h light/dark cycle and were allowed free access to water and feed. All animal experiments were approved by the Laboratory Animal Welfare and Ethics Committee of Third Military Medical University (No. AMUWEC20182139).

### Tamoxifen injection

To induce tdTomato expression, tamoxifen (Sigma-Aldrich, St. Louis, MO, USA; T5648) dissolved in corn oil was injected intraperitoneally at 120 mg/kg body weight into MG lineage-tracing mice at P21 every day for 5 days, followed by a 7-day suspension^42^.

### Samples preparation and immunofluorescence (IF) staining

As described previously^19,82^, the eyes were fixed with 4% paraformaldehyde for 2 h after removing cornea and lens. Then, the eye cups were dehydrated in 30% sucrose solution overnight, followed by embedding using optimum cutting temperature compound and quick freezing at −80 °C. The samples were cut into 12 μm-thick slices via a cold microtome (Leica, Germany) and preserved at −20 °C. Cell climbing slices were washed with PBS for three times and fixed with 4% paraformaldehyde for 15 min at room temperature. For immunostaining, sections were washed with 0.01 M PBS three times and blocked with 0.3% Triton X-100 containing 3% bovine serum albumin (BSA) for 30 min. The slices were incubated with primary antibodies at 4 °C overnight, followed by secondary antibodies at 37 ° C for 1 hour. The slices were photographed via confocal laser scanning microscopes (Zeiss LSM 880 and Nikon AX) after nuclei were counterstained with DAPI (Beyotime, C1005). The antibodies and dilutions used for IF were showed in Table 1.

**Table 1.** Antibodies used for immunofluorescence staining and western blotting (WB)

| Name | Host | Supplier | catalogue | Application |
| --- | --- | --- | --- | --- |
| Primary antibodies |  |  |  |  |
| GFAP | rabbit | Dako | Z0334 | IF (1:500) |
| GS | mouse | Sigma | MAB302 | IF (1:500) |
| Sox9 | rabbit | Abcam | ab185966 | IF (1:500) |
| Pax6 | mouse | BioLegend | 862002 | IF (1:300) |
| Chx10 | mouse | Santa Cruz | sc-374151 | IF (1:300) |
| Chx10 | sheep | Sigma | AB9016 | IF (1:300) |
| cone-arrestin | rabbit | Sigma | AB15282 | IF (1:500) |
| Rhodopsin | mouse | Abcam | ab5417 | IF (1:500) |
| Sox2 | rabbit | Abcam | ab97959 | IF (1:500) |
| Lamp1 | rabbit | Abcam | ab24170 | IF (1:400) |
| CSPGs | mouse | Sigma | C8035 | IF (1:300) |
| Collagen-1 | rabbit | Abcam | ab34710 | IF (1:500) |
| Mmp2 | rabbit | Abcam | ab92536 | IF (1:400) |
| Mmp9 | rabbit | Abcam | ab38898 | IF (1:400) |
| PKCa | mouse | Abcam | ab11723 | IF (1:500) |
| HuC/D | mouse | Invitrogen | A21272 | IF (1:300) |
| Vimentin | mouse | Santa Cruz | sc-6260 | IF (1:300) |
| Ki67 | rabbit | Abcam | ab16667 | IF (1:500) |
| HuNu | mouse | Abcam | ab191181 | IF (1:200) |
| GFAP | rabbit | Abcam | ab7260 | WB (1:1000) |
| Mmp2 | rabbit | Abcam | ab92536 | WB (1:1000) |
| β-Actin | rabbit | Abclone | AC038 | WB (1:10000) |
| **Secondary antibodies** |  |  |  |  |
| anti-rabbit IgG Alexa-Fluor-647 | goat | Invitrogen | A-21245 | IF (1:1000) |
| anti-mouse IgG Alexa-Fluor-647 | goat | Invitrogen | A-21235 | IF (1:1000) |
| anti-rabbit IgG Alexa-Fluor-488 | goat | Invitrogen | A-11008 | IF (1:1000) |
| anti-mouse IgG Alexa-Fluor-488 | goat | Invitrogen | A-11001 | IF (1:1000) |
| anti- sheep IgG Alexa-Fluor-488 | donkey | Invitrogen | A-11005 | IF (1:1000) |
| anti-rabbit IgG Alexa-Fluor-568 | goat | Invitrogen | A-11036 | IF (1:1000) |
| anti-mouse IgG Alexa-Fluor-568 | goat | A-11031 | Invitrogen | IF (1:1000) |
| anti-rabbit IgG(H+L) HRP | goat | AS014 | Abclone | 1:5000 |

### The quantity of MG number

To count the numbers of MG and migrated MG in the retinas of C57, PBS and hAECs groups, the number of Sox9 positive cells in the middle of the temporal side of the retina with 159 μm × 159 μm were counted for statistical analysis.

### scRNA-seq analysis

The retinas were digested into single-cell suspensions using GEXSCOPE^TM^ Tissue Dissociation Solution (Singleron, China), and tdTomato^+^ cells were isolated for scRNA-seq using fluorescence-activated cell sorting. Subsequently, the cell suspension was loaded onto microfluidic chips (Singleron), and scRNA-seq libraries were constructed using a GEXSCOPER Single-Cell RNA Library Kit (Singleron) according to the manufacturer’s instructions. Sequencing was performed using an Illumina 6000 (Illumina, USA) with 150-bp paired-end reads. The Cell Ranger pipeline was used to process the raw data, and the original counts were inputted into the R platform. The Seurat package integrates different data, followed by an analysis. Cells with a low number of detected genes (<200) or a high percentage of detected mitochondrial genes (>20%) were considered low-quality and removed. We normalized, scaled the expressions of each cell and find 2000 variable features using NormalizeData, FindVariableFeatures and ScaleData functions in Seurat package. The RunPCA function performed principal component analysis (PCA) dimensionality reduction and RunHarmony function in Harmony package was used to integrate different databases and correct batch effects across samples^36^. RunUMAP function assessed UMAP dimension reduction, clustering after FindNeighbors and FindClusters. The FindAllMarkers function calculated the marker genes of clusters (genes with *p*_val_adj < 0.05 and avg_log2FC > 0.25). The AUCell package was used to calculate the RPC scores. Pseudotime analysis was performed using the monocle3 package^83^.

### Correlation analysis between developmental and adult retinas

The average expressions of cells in each celltype were calculated using AverageExpression function and were used to calculate correlation coefficients between developmental and adult retinas using cor function.

### ROGUE analysis for cell purity

ROGUE package, an entropy-based method was used to quantify the purity of identified MG subclusters accurately and robustly ^84^.

### Pseudobulk analysis

To avoid false discoveries in differential expression analysis between different groups in single-cell data, we performed differential state analysis to identify DEGs between healthy and degenerative retinas based on aggregated pseudobulk data using muscat package ^85^. Genes (*P* < 0.05, adjusted *P* value < 0.1 and log2foldchange > 0.5 for 3 wpn and *P* < 0.05, adjusted *P* value < 0.05 and log2foldchange > 0.5 for 8 wpn) were recognized as DEGs.

### Enrichment analysis

GO and KEGG enrichment analysis was performed using enrichGO and enrichKEGG functions in clusterProfiler package. GO and KEGG terms with adjusted *P* values < 0.05 were considered significantly enriched. GSEA was conducted using gseGO function in clusterProfiler package and GSEA terms with *P* value < 0.05, adjusted *P* value < 0.25 and NES > 1 were statistical significantly.

### Cell-labeling and subretinal transplantation of hAECs

hAECs were purchased from Shanghai iCELL Biotechnology and were cultured in Dulbecco’s modified Eagle medium (DMEM) containing 10% fetal bovine serum (FBS), 2 mM L-glutamine, 55 μM of 2-mercaptoethanol, 1% nonessential amino acid, 1% antibiotic-antimycotic 1 mM of sodium pyruvate, (Gibco, Carlsbad, CA) and 10 ng/mL EGF (Peprotech)^86^. When confluency reached 50%, hAECs were transfected with lentivirus-carrying CMV promoter-driven EGFP for 16 h and replaced with the fresh medium. We used the EGFP-labeled hAECs for subretinal transplantation after cell passaging. As previously reported, hAECs were digested with 0.25% trypsin supplemental with 0.02% EDTA (Solarbio, 9002-07-7) and resuspended in PBS (Hyclone, SH30256.01) supplementing with 0.005% DNase I (STEMCELL Technologies, 07900) at a concentration of 1×10^5^ cells/μl. rd10 mice were anesthetized at 2 weeks postnatal (2 pwn) by intraperitoneal injection of 1% sodium pentobarbital (2.5ml/kg body weight) and used 1% tropicamide for pupil dilation. 1 μl hAECs suspension were injected into subretinal space using a 33-gauge Hamilton needle (Hamilton) and an equal volume of PBS was used as the control. All mice received oral cyclosporine A (210 mg/l, Sandoz, Camberley) dissolved in drinking water from 24 h before the operation to 2w post-transplantation.

### Western blotting

Total Protein Extraction Kit for WB (Immunoway, RS0024) was used to extract protein from the whole retina, and protein concentrations were determined using a BCA assay kit (Beyotime, P0010). An equal amount of protein samples was loaded on a 12% SDS-PAGE gel and electroblotted to a PVDF membrane. Then, the membranes were blocked with Y-Tec 5min and Ready-to-Use Blocking Buffer (Yoche, YWB0501) 20 min and incubated with primary antibodies overnight at 4 °C. On the next day, membranes were incubated with secondary antibody. After detection with Pierce™ ECL Western Blotting Substrate (Thermo, 32106), we scanned proteins on the membranes with a Bio-Rad exposure system (BioRad). The antibodies and dilutions used for WB were showed in Table 1.

### Electroretinogram (ERG) recording

As described previously^19,82^, animals were darkly adapted for at least 12 hours before scotopic ERG recording. The gold electrode rings contacted the cornea as the recording electrodes after anesthetizing mice. The reference electrodes were placed into the mice’s mouths and the ground electrodes clamped the mice’s tails. We recorded at different light intensities (0.001, 0.01, 0.1 1.0, and 3.0 cd·s/m^2^) using an ERG recording device (MAYO, Japan) and used GraphPad Prism 9.2.0 software to analyze the data.

### Light/dark transition test

The visual behavior test evaluated the visual sensitivity as previously reported^87^. The test device contained a dark box (30 cm × 20 cm), a light box (30 cm × 30 cm), and a hole (10 cm × 10 cm) between two boxes for mice shuttling. Mice received more than 12 hours of dark adaptation and stayed in the dark box for 5 min before recording started. Then, turn on the lamp to maintain a 400 lux light intensity in the light box, and the mice can shuttle freely between the dark and light box for 5 min.

### In vivo fluorescence fundus imaging and optical coherence tomography

A standard confocal laser scanning ophthalmoscope (Phoenix Contact, Germany) was used to image the retinas of mice and EGFP-labeled hAECs. We anesthetized the mice and dilated their eyes, and the microscope lens contacted the eye perpendicularly at the central cornea to acquire the fundus and OCT images.

### Co-culture of MIO-M1 cells and hAECs

MIO-M1, human Müller glia line, was kindly gifted by Professor Jinfa Zhang of Shanghai Jiao Tong University, Shanghai, China. MIO-M1 was cultured in high glucose DMEM supplemented 10% FBS and 1% Penicillin-Streptomycin solution. When the cell confluence reached 80%-90%, they were digested with trypsin and passaged. All cultured cells are incubated in a standard incubator (37°C, 5%CO2, saturated humidity). According to our previous study^19^, MIO-M1 cells were seeded on cell culture plates at a density of 2 × 10^4^ cells/cm^2^ culture area, and the same quantity of hAECs were seeded on cell culture inserts with 0.4-μm aperture membranes (Corning, USA). MIO-M1 cells and hAECs were co-cultured for 72 h for the following analysis.

### Cell cycle assay

According to the manufacturer’s instructions, a cell cycle test was performed using DNA Content Quantitation Assay (Cell Cycle) (Solarbio, CA1510). Briefly, after being digested and centrifuged, cells were fixed with 70% pre-cold ethanol overnight, and cell precipitates were incubated with 100µl RNase A solution in a water bath at 37 °C for 30 min, followed by incubation with 400µl PI solution at 4 °C for 30 min (protect from light). Then, flow cytometry was performed using BD FACS Aria II flow cytometer for detection.

### Cell migration assay

Cell migration assay was performed using transwell plates with 8-μm aperture membranes (Corning, USA)^21^. Before coculturing, MIO-M1 cells were treated with 1 μg/ml Mitomycin C (MedChemExpress, HY-13316) for one hour to block cell proliferation. hAECs were seeded in the lower chamber, and MIO-M1 cells were in the upper chamber followed by coculturing for 72 h. The migrated MIO-M1 cells were stained with a Crystal violet solution (Solarbio, C8470).

### Statistical analysis

Data from more than three biological replicates were analyzed and presented as mean ± SD/SEM. Students’ *t* test was performed for single comparison between two groups, multiple *t* tests with Benjamini-Hochberg correction were performed for multiple comparisons between two groups, and one-way analysis of variance (ANOVA) followed by Tukey’s multiple tests was performed for comparisons between multiple groups via GraphPad Prism 9.2.0. Differences were accepted as significant at *p* < 0.05.

### Data Availability

Data and code of scRNA-seq will be made available upon required. Other data can be acquired in the main txt and supplemental files

## Acknowledgments

This study was supported by funding from the National Natural Science Foundation of China (No. 31930068, 82271104), the National Key Research and Development Program of China (Grant No. 2021YFA1101203).

## Author Contributions

H.X. conceived, designed, and supervised the project, revised manuscript; X.F. and J.X. supervised the project, revised manuscript; H.Gao. performed experiments, analyzed data and wrote the manuscript; Z.Y. assisted with experiments and revised manuscript; Y.Z., X.H., Z.C. and X.C. assisted with in vivo experiments; L.G., J.K., X.L., assisted with in vitro experiments; T.Z. provided guidance for this project; and H.Gong. revised manuscript.

## Declaration of Interests

The authors declare that they have no conflicts of interest.

## Notes

### Competing Interest Statement

The authors have declared no competing interest.

