## Supplemental Figures for "Human Amniotic Epithelial Cells Promote Chx10^−^/Pax6^+^ Müller Glia Subpopulation Reprogramming into Photoreceptor-like Cells"

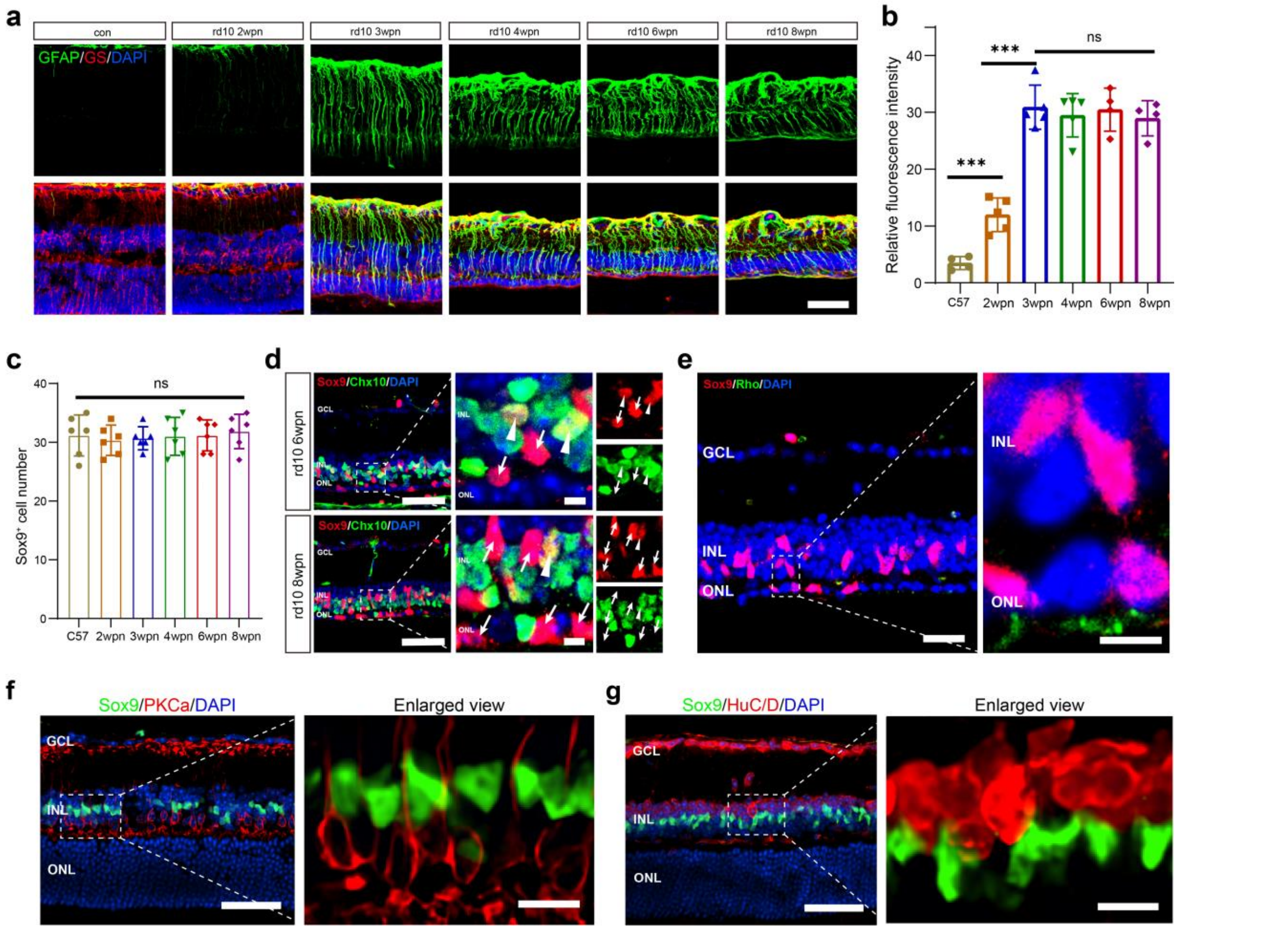


**Figure S1.** Müller glia increased the gliosis level and didn't express photoreceptor marker in the degenerative retina.

**a** Representative fluorescence images of GFAP and GS staining in the retinas of C57 and rd10 mice from 2 wpn to 8 wpn.

**b** relative quantitative analysis of GFAP staining.

**c** The number of $Sox9^+$ nuclei in the retinas of C57 and rd10 mice from 2 weeks postnatal to 8 wpn.

**d** Representative immunofluorescence images of Sox9 and Chx10 in the retinas of rd10 mice at 6 wpn and 8 wpn, arrow: $Sox9^+Chx10^-$ nuclei, arrowhead: $Sox9^+Chx10^+$ nuclei.

**e** Representative fluorescence images of Sox9 and rhodopsin in the retina of rd10 mice at 8 wpn.

**f** Representative fluorescence images of Sox9 and PKCa.

**g** Representative fluorescence images of Sox9 and HuC/D.

Data are presented as mean ± SD, n = 6 eyes per group. For statistical analysis, one-way ANOVA followed by Tukey's multiple tests (b, c) is applied. * $p < 0.05$, ** $p < 0.01$, *** $p < 0.001$; ns, no significance. Scale bars, 50 μm (a, d, f, g), 20 μm (e), 10 μm (enlarged image of f, g), 5 μm (enlarged images of d, e).

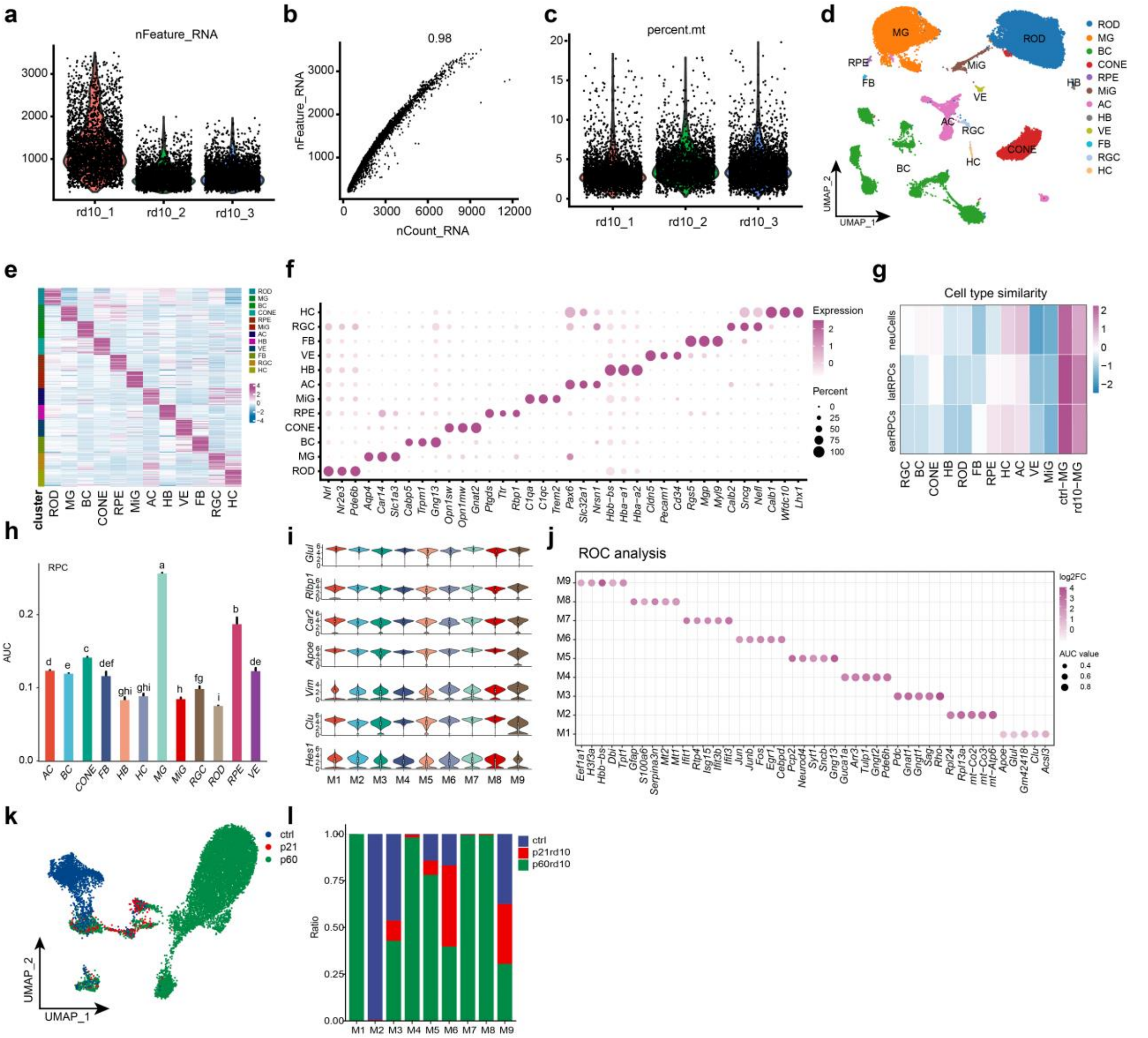


**Figure S2.** scRNA-seq analysis of Müller glia (MG) in the retina.

**a-c** The quality control analysis of scRNA-seq data.

**d** UMAP plot showing unsupervised clustering of major cell types in the retina.

**e** Heatmap of top 30 differentially expressed genes of each cell type compared to all other cell types.

**f** heatmap showing the correlations between the cell types from the adult and development retinas

**g** Dotplot showing expression patterns of major retinal class markers.

**h** Barplot showing the RPC score of each cell type using the AUCell package.

**i** Violin plot showing the expressions of classical MG marker genes in MG subtypes.

**j** Dotplot showing marker genes of MG subtypes by ROC analysis

**k** UMAP plot showing MG from different group.

**l** Fractions of cells (dot size) in each MG subtype from different group.

earRPCs: early retinal progenitor cells; latRPCs: late retinal progenitor cells; neuCells: neurogenic cells

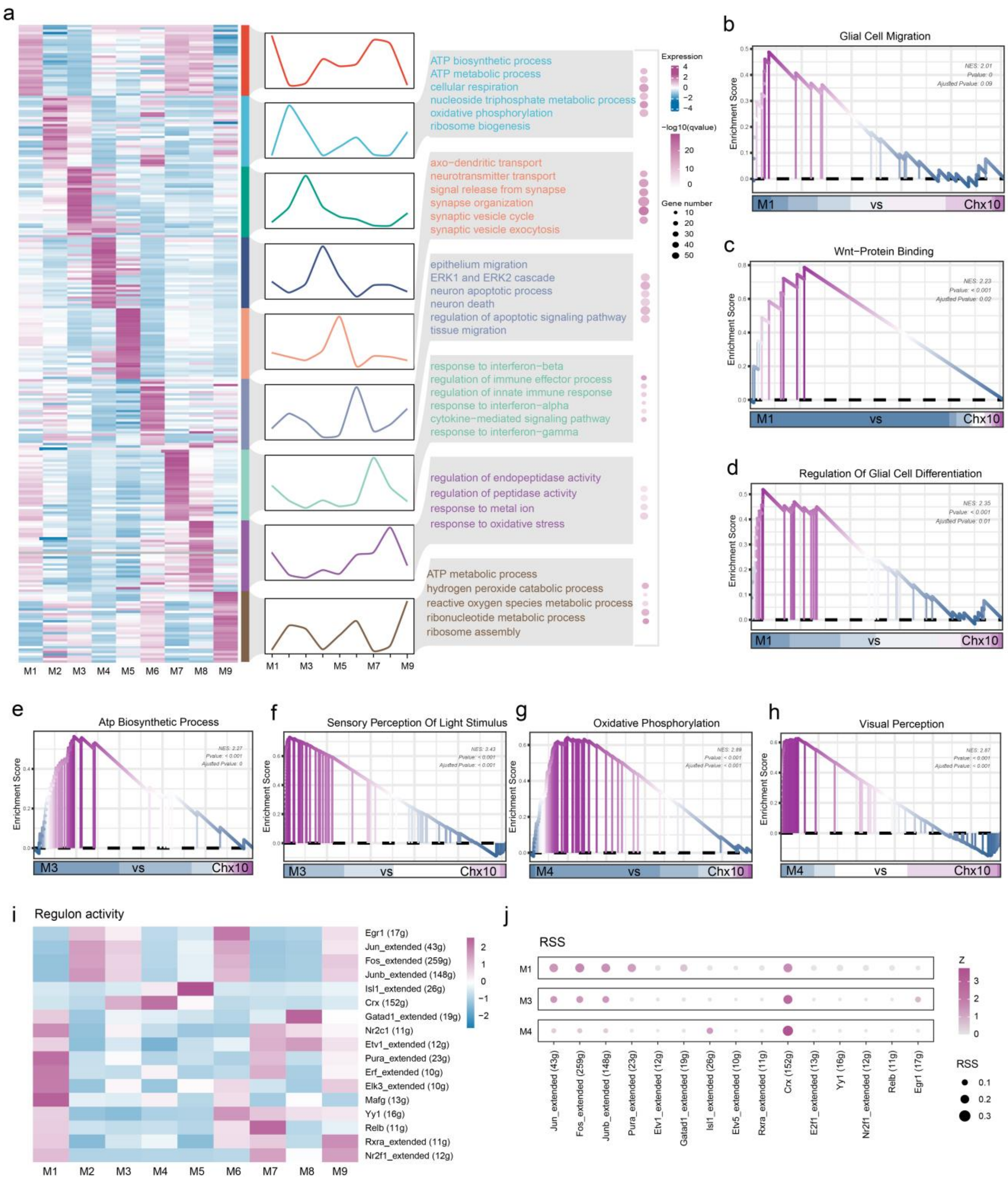


**Figure S3.** Enrichment analysis of MG subclusters.

**a** Representative GO terms of marker genes of different MG subclusters.

**b-d** GSEA plot showing glia cell migration, Wnt-protein binding, regulation of glial cell differentiation in M1 and Chx10$^+$ subclusters (M5 and M6).

**e, f** GSEA plot showing ATP biosynthetic process and sensory perception of light stimulus in M3 and Chx10$^+$ subclusters (M5 and M6).

**g, h** GSEA plot showing oxidative phosphorylation and visual perception in M4 and Chx10$^+$ subclusters (M5 and M6).

**i, j** GSEA plot showing oxidative phosphorylation and visual perception in M4 and $Chx10^+$ subclusters (M5 and M6).

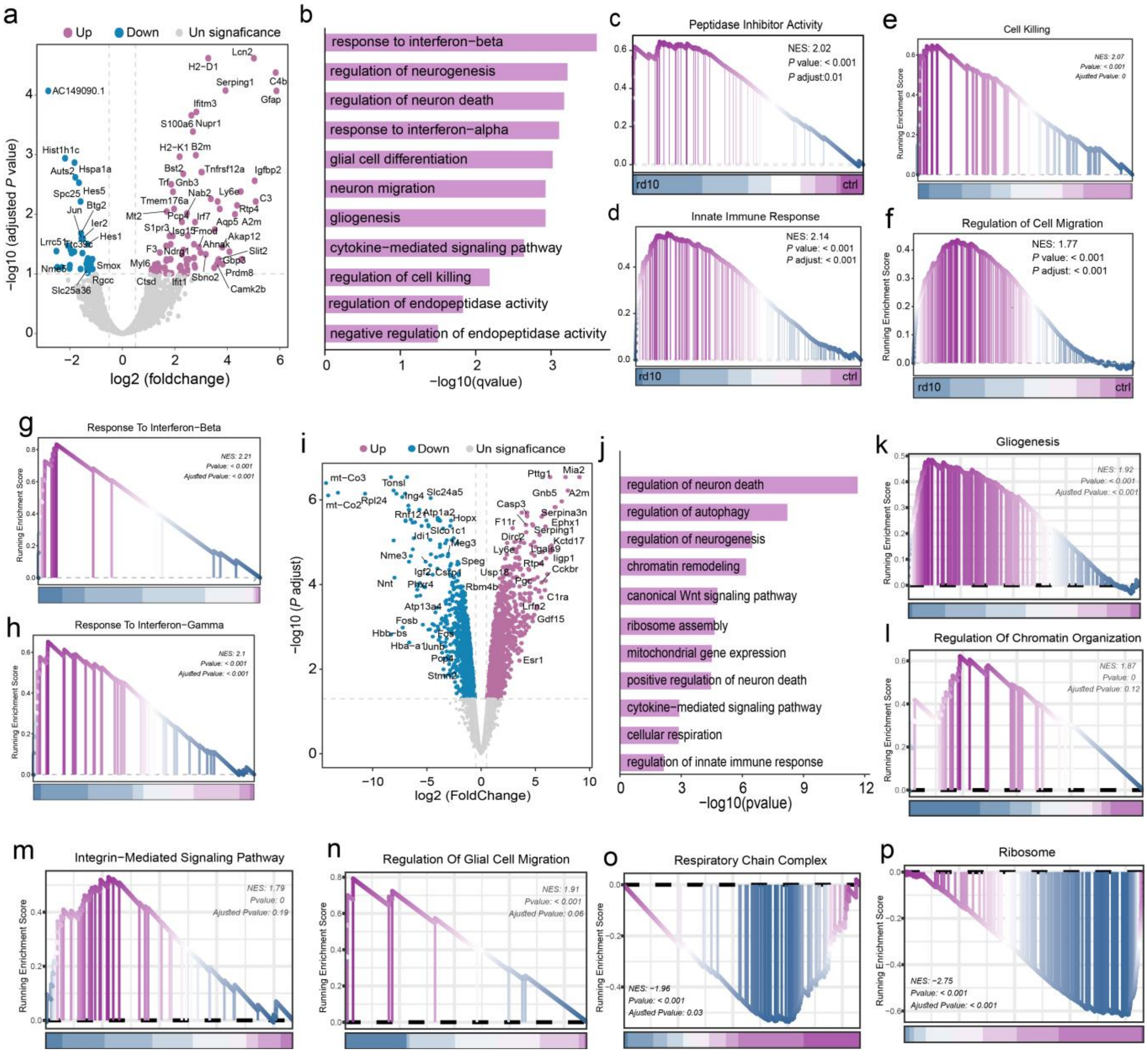

**Figure S4.** differential expression analysis between healthy retina and degenerative retina.

**a** Volcano plot showing DEGs between MG from healthy and degenerative retinas at 3 wpn.

**b** GO enrichment terms of DEGs between MG from healthy and degenerative retinas at 3 wpn.

**c-h** GSEA plot showing peptidase inhibitor activity, innate immune response, cell killing, regulation of cell migration, response to interferon-beta/gamma in MG from healthy and degenerative retinas at 3 wpn.

**i** Volcano plot showing DEGs between MG from healthy and degenerative retinas at 8 wpn.

**j** GO enrichment terms of DEGs between MG from healthy and degenerative retinas at 8 wpn.

**k-p** GSEA plot showing gliogenesis, regulation of chromatin organization, integrin-mediated signaling pathway, regulation of glia cell migration, respiratory chain complex and ribosome in MG from healthy and degenerative retinas at 8 wpn.

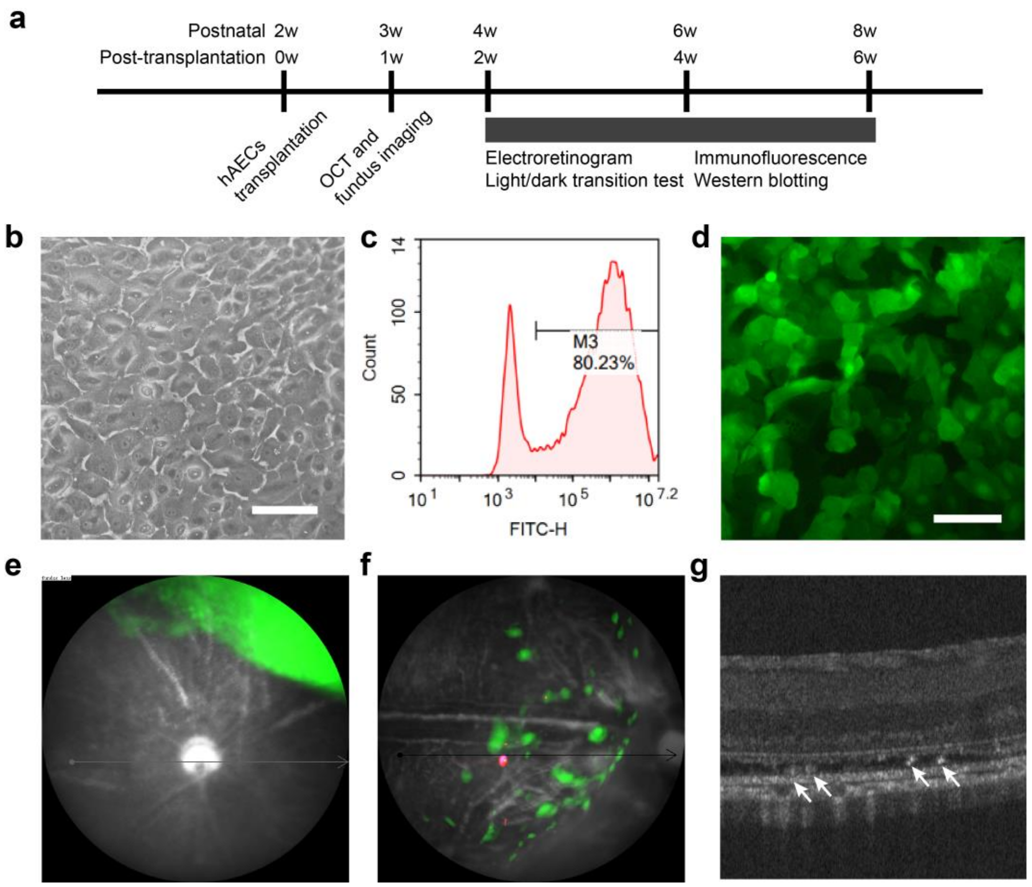


**Figure S5.** The retina sub-space transplantation of hAECs.

**a** Experimental design.

**b** Representative image of hAECs morphology. Scale bars, 100 µm.

**c** FACS analysis of EGFP-positive hAECs.

**d** Representative image of EGFP-labeled hAECs infected with lentivirus. Scale bars, 100 µm.

**e, f** In vivo fluorescence fundus images showing the distribution of EGFP-labeled hAECs in retina just after transplantation (e) and at 1 week after transplantation (f).

**g** OCT image of retina with transplanted hAECs.

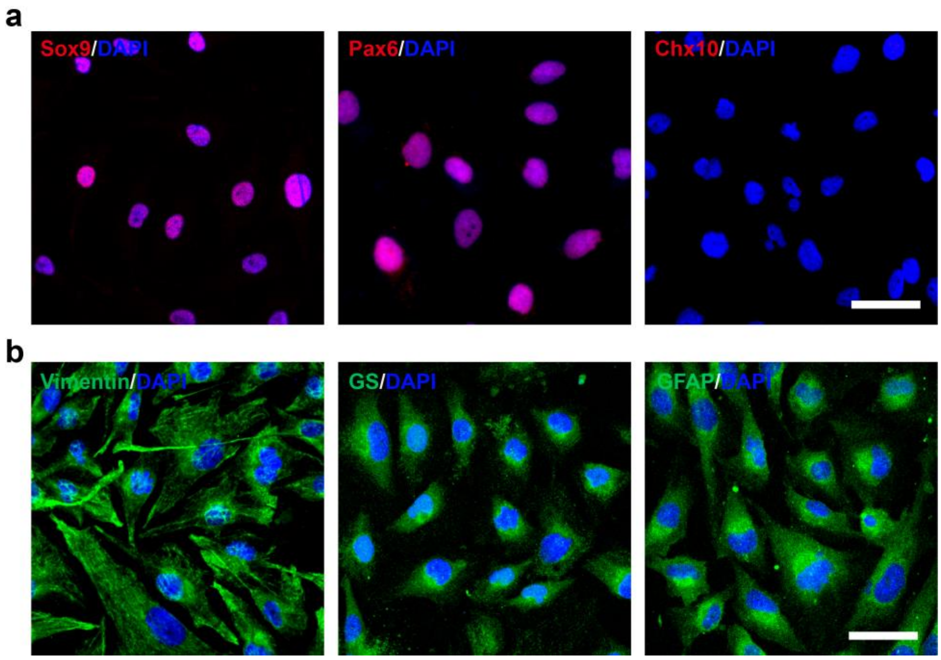


**Figure S6.** The identification of human Müller glia line MIO-M1.

**a** Representative fluorescence images of nuclear antigens Sox9, Pax6 and Chx10.

**b** Representative fluorescence images of cytoplasmic antigens Vimentin, GS and GFAP.

Scale bars, 50 μm.

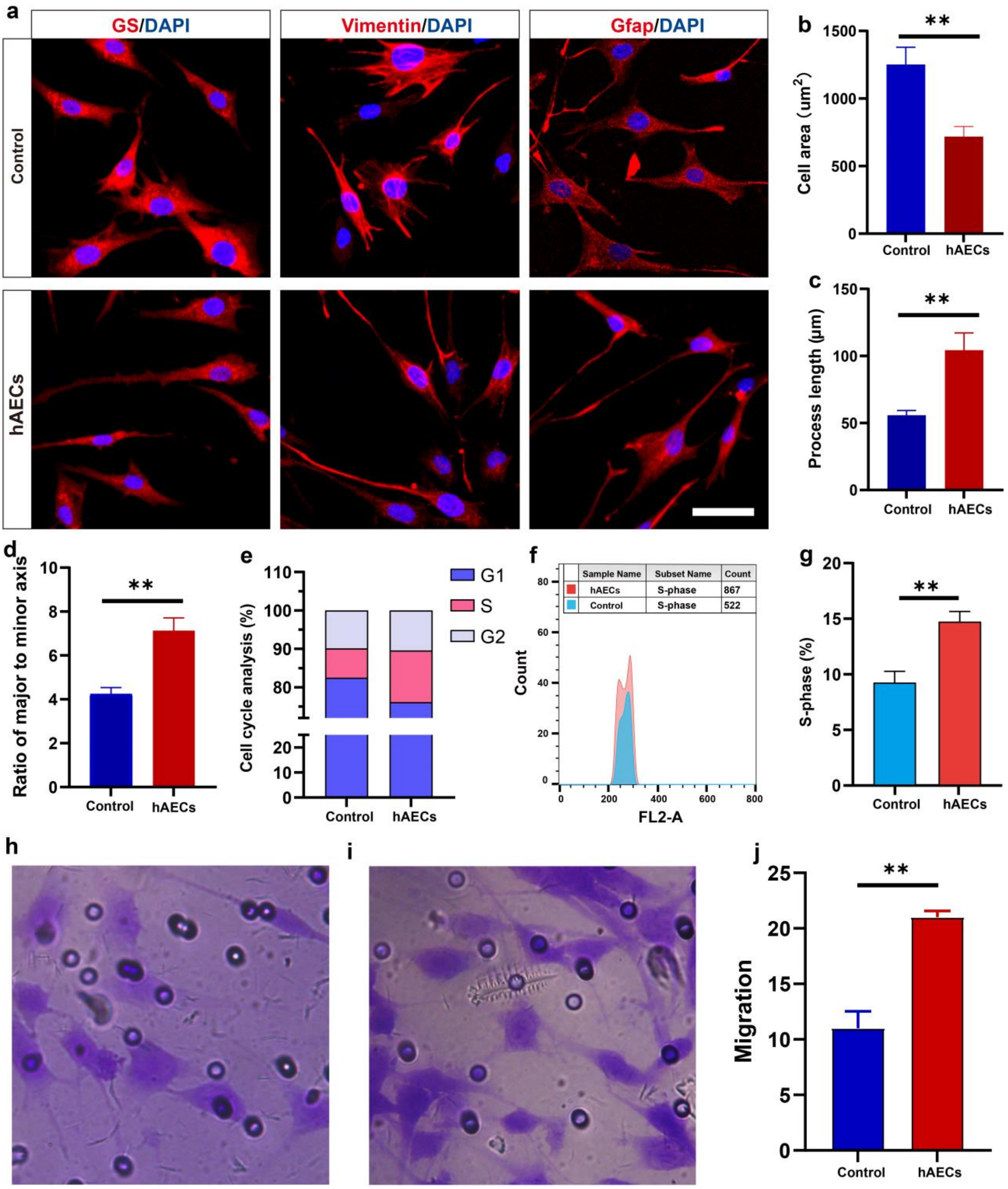


**Figure S7.** hAECs promote the proliferation and migration of MG in vitro.

**a** Representative fluorescence images of GS, Gfap, and Vimentin in control and hAECs groups.

**b-d** Quantitative analysis of morphology in terms of cell area (B), process length (C), and the ratio of major to minor axis (D).

**e** Percentage of cells in the G1 phase, S phase, and G2 phase.

**f** Representative flow plots of cells in the S phase in control and hAECs groups.

**g** Statistical analysis of the percentage of cells in the S phase.

**h, i** Representative crystal violet staining images of migration assay in control (H) and hAECs (I) groups.

**j** The number of cells migrated in control and hAECs groups.

Data are presented as mean ± SD, n = 6 (b-d, g), n = 3 (j). For statistical analysis, student *t* test is applied. * $p < 0.05$, ** $p < 0.01$, *** $p < 0.001$; ns, no significance.

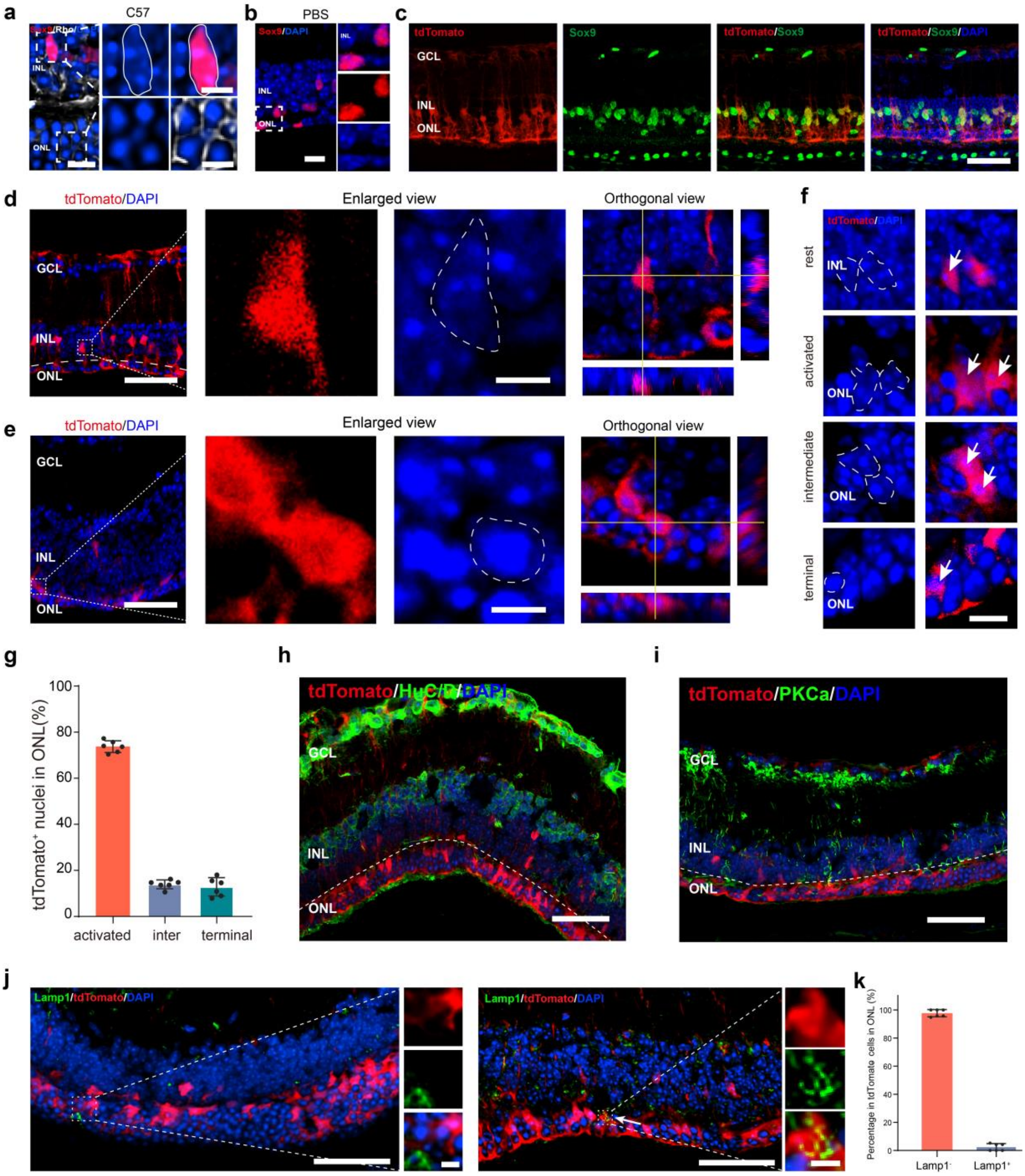


**Figure S8.** hAECs promote MG reprogramming into photoreceptor-like cells rather than other neurons.

**a, b** Representative images of Sox9 staining and nuclear architecture at 6 wpt in the C57 (a) and PBS (b) groups.

**c** Representative images of tdTomato and Sox9 staining in MG lineage-tracing mice with retinal degeneration.

**d, e** Representative images of the architecture of tdTomato$^{+}$ nuclear in PBS (d) and hAECs (e) groups.

**f** Representative images of MG-derived photoreceptor-like nuclei transition through rest, initial, intermediate and terminal stages.

**g** Quantitative analysis of MG-derived photoreceptor-like nuclei in the outer nuclear layer at 6 wpt.

**h** Representative image of tdTomato and HuC/D staining at 6 weeks after hAECs transplantation.

**i** Representative image of tdTomato and PKCa staining at 6 weeks after hAECs transplantation.

**j, k** Representative images of tdTomato and Lamp1 staining at 6 wpt. Arrow indicates the co-labeling of tdTomato and Lamp1.

Data are presented as mean ± SD, n = 6 eyes per group. Scale bars, 50 μm (c, d, e, h, i, j), 10 μm (a, b), 5 μm (enlarged images of a, d, e, j; f).

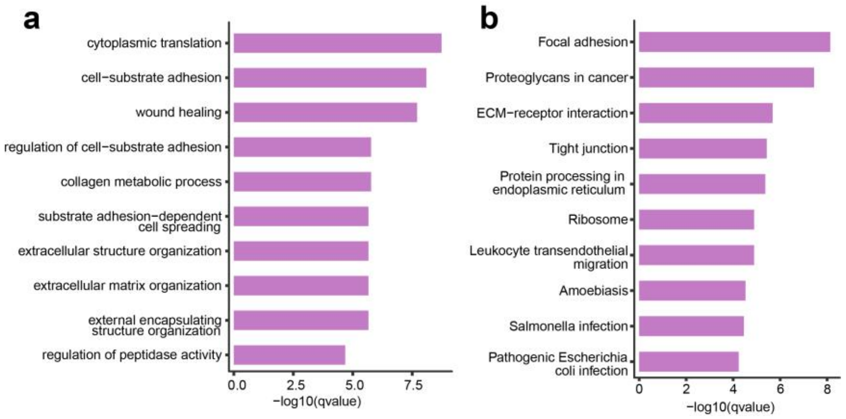


**Fig S9.** Functional analysis of top 200 genes in hAECs.

**a** The top 10 terms of GO (biological process) analysis.

**b** The top 10 pathways of KEGG analysis.

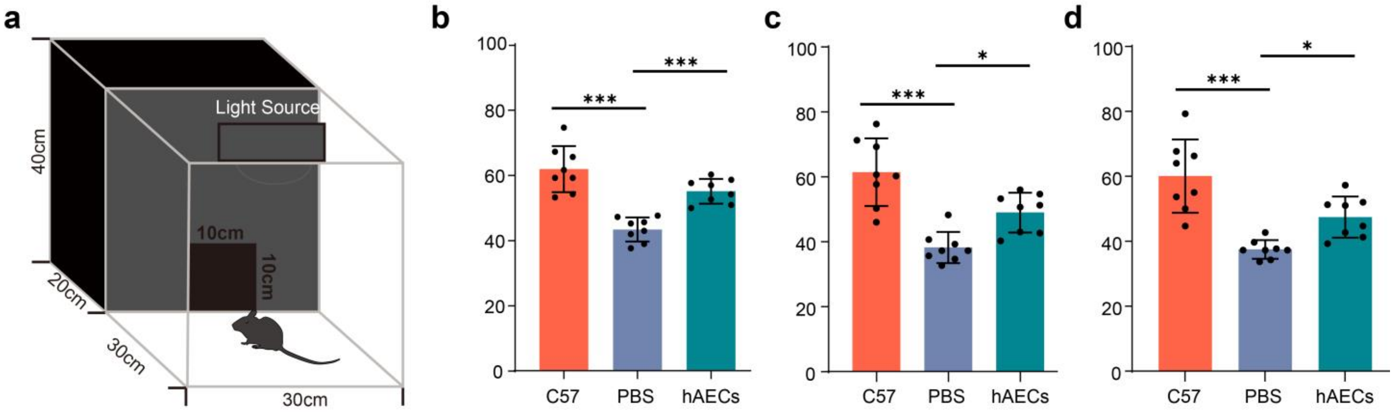


**Figure S10.** hAECs transplantation improved the visual behaviors of rd10 mice.

**a** Diagram showing the device of light-dark transition test.

**b-d** Statistical analysis of percentage of time the mice spent in dark box in C57, PBS and hAECs groups at 2 wpt (b), 4 wpt (c) and 6 wpt (d).

Data are presented as mean ± SD, n = 8 mice per group. For statistical analysis, one-way ANOVA (b, c, d) followed by Tukey's multiple tests is applied. * $p < 0.05$, ** $p < 0.01$, *** $p < 0.001$; ns, no significance.

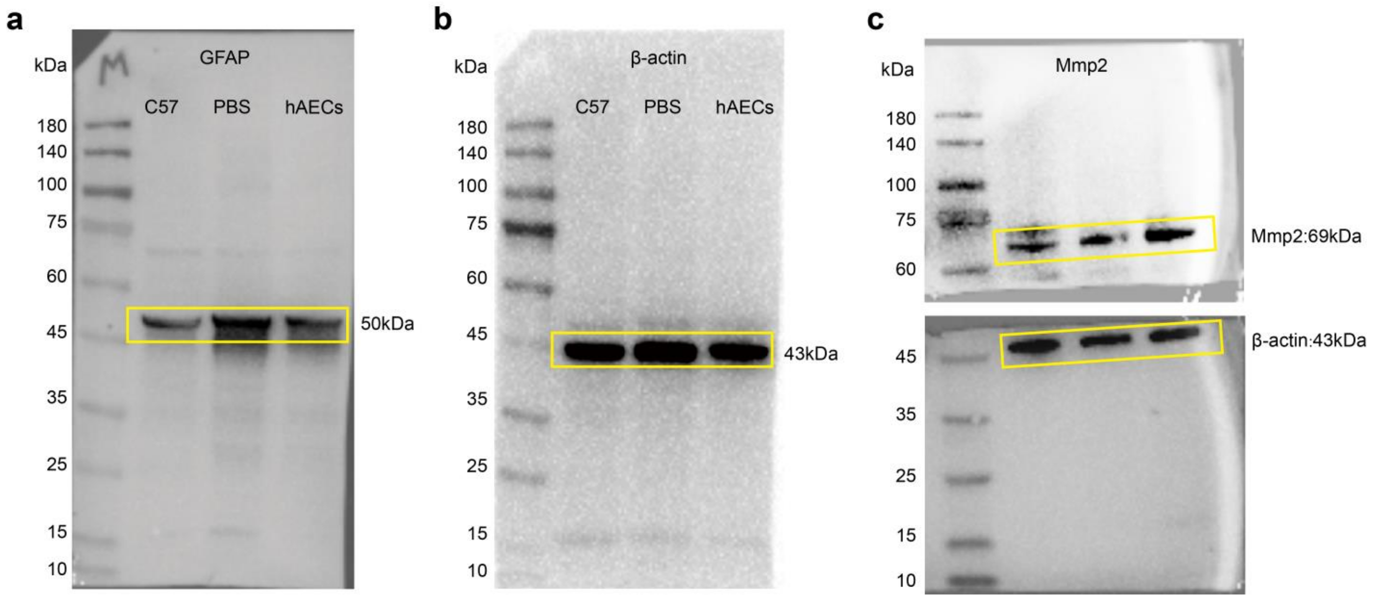


**Fig S11.** The whole western blot images.
**a-b** are affiliated to Fig 5c.
**c** is affiliated to Fig 5i.
